# Whole genome similarity provides a rapid, robust framework for classification of fungal taxa from the genus rank to intraspecies variants

**DOI:** 10.64898/2026.09.04.746300

**Authors:** Hayden Johnson, Boris A. Vinatzer, Reza Mazloom, Kassaye Belay, Niklaus J Grünwald, Jessie K. Uehling

**Affiliations:** Department of Botany and Plant Pathology 2701 NW Orchard Ave. Corvallis OR 97330, USA; School of Plant and Environmental Sciences, Virginia Tech, Blacksburg, VA 24061, USA; Department of Computer Science, Virginia Tech, Blacksburg, VA 24061, USA; Graduate Program in Genetics, Bioinformatics, and Computational Biology, Virginia Tech, Blacksburg, VA 24061, USA; Horticultural Crops Disease and Pest Management Research Unit, USDA ARS, 3420 NW Orchard Ave., Corvallis, OR 97330, USA

**Keywords:** whole-genome similarity, fungal genomics, oomycete genomics, sourmash, fungal taxonomy, ANI, average nucleotide identity, systematics, diversity, evolution

## Abstract

Rapid and accurate microbial identification is critical for interpreting biological data in basic research and when making applied decisions on how to effectively treat patients and control human, animal, and plant diseases. Advancements in high-throughput sequencing have the potential to expedite fungal species identification and thus fungal biological research; however, analyses of ever-increasing numbers of genomes also present computational challenges. In this work, we evaluated how whole-genome similarity can serve as the basis for accurate classification and identification across the Kingdom Fungi and the Phylum Oomycota. Results from the computationally efficient k-mer–based tool sourmash are compared with those from more computationally demanding BLAST-based similarity method ANIb, as well as with conventional phylogenomic approaches, including maximum-likelihood concatenated ortholog trees and SNP-based methods. We observed that sourmash delivers orders-of-magnitude gains in speed and memory efficiency while maintaining strong concordance with phylogenomic methods. We found that a k-mer size of 21 is robust for genus and species determination, and that larger k-mers are well-suited for identification below the species rank. These results demonstrate that k-mer-based whole-genome similarity provides a scalable framework for fungal classification, enabling rapid analysis, lowering computational and bioinformatic barriers, and supporting the development of efficient identification pipelines.

## Introduction

Taxonomically grouping microbes into genetically and phenotypically discrete classes is fundamental to both scientific research and societal applications. Identification of an unknown microbe as a member of such a class enables reliable prediction of that organism’s characteristics and thus informs appropriate responses - for example, selecting an effective therapeutic agent for a microbial infection. In fungal taxonomy, species are traditionally treated as the primary unit of classification, but many morphologically plastic fungi harbor cryptic species that may differ in medically or agriculturally important traits [1, 2]. Another challenge is that historically, taxonomic assignments for fungi and oomycetes have relied on morphological, physiological, and sexual compatibility characteristics. Although these approaches have provided a strong foundation for classification, their utility is limited, for many lack distinctive morphological features, are challenging to culture in lab, and have elusive reproductive dynamics, making robust identification challenging [3].

More recently, single- and multi-gene phylogenetic analyses have been employed to delimit species. However, these approaches often lack the resolution needed to unambiguously discriminate among closely related species and, even more so, among intraspecific lineages [4]. Today, sequence-based methods have progressed in scope and resolution beyond single and multi-gene sequencing to whole-genome based approaches, which enable single-copy ortholog gene [5, 6] or SNP-based phylogenies to resolve taxa that barcoding or multi-locus approaches cannot. However, these workflows require access to substantial computational resources, significant time, and bioinformatics expertise, especially for analyses at large scales. The accelerating production and availability of fungal genome sequences highlight the need for more efficient methods to rapidly infer fungal identity from these data.

Whole genome similarity-based species delimitation [3, 7] is widely used in prokaryotes and is generally computed as average nucleotide identity (ANI) which initially served as a computational proxy for DNA–DNA hybridization [8]. While ANI is well established in prokaryotic taxonomy, its use in fungi is relatively recent but increasing steadily [9–25]. It is largely focused on yeasts, molds, and plant pathogens, with no consensus on a preferred approach (Supplementary Table 1). This lack of consensus is compounded by the fact that ANI lacks a single formal definition and is calculated using diverse methodologies which vary in their underlying assumptions and computational frameworks, underscoring the urgent need to establish standardized benchmarks for employing whole-genome sequence data in fungal classification.

ANI estimation methods range from classic alignment-based approaches such as ANIb, the legacy gold standard method based on BLAST [26], OrthoANI [27], OrthoANIu [28], ANIm [28, 29], and single copy ortholog-focused [22] ANI methods, which estimate mean identity across aligned genomic regions. In contrast, faster approaches such as FastANI [7] approximate sequence mapping in place of alignment. Even faster alignment-free, k-mer–based sketching methods (e.g., used by Mash [30], Dashing [31], GSearch [32], and sourmash [33]) estimate ANI from genome-wide k-mer similarity using variations of MinHash, HyperLogLog, or related techniques. Sketching approaches are distinct from the alignment and mapping methods in that they leverage information throughout the entire genome, rather than considering only alignable or orthologous regions. Critically, these methods create compact ‘fingerprints’ of genomes that require far less memory and permit extremely fast pairwise comparisons and database searches, enabling scalable analysis as reference databases grow to hundreds of thousands of genomes and beyond. This scalability is particularly relevant for fungi, whose genomes are generally larger and contain more repetitive sequences than those of prokaryotes, making alignment-based ANI increasingly computationally prohibitive at scale.

To address this need for rapid, high-resolution identification, we chose to comprehensively evaluate the utility of the sketching tool sourmash for fungal and oomycete classification. Like other sketching methods, sourmash is characterized by high speed and low memory requirements compared to non-sketch-based methods. Sourmash is distinguished from other sketching methods by its deployment of FracMinHash sketches which retain a fixed fraction of k-mers rather than a fixed sketch size, resulting in sketches that scale proportionally with genome size [34]. FracMinHash sketches not only allow for Jaccard similarity estimation but also permit accurate max containment estimation, a metric defined by the proportion of shared unique k-mers relative to the number of unique k-mers in the smaller genome. Sourmash estimates ANI from metrics such as Jaccard similarity and max containment by applying a theoretical conversion assuming a simple fixed rate mutation model [35]. Further, sourmash is extensible to large, indexed databases and metagenomic analyses.

To explore if whole genome ANI could provide a robust framework for classification of fungal taxa from the genus rank to infraspecific variants here we (1) applied sketch-based, sourmash-derived ANI to fungi and oomycetes, (2) evaluated both Jaccard- and max-containment similarity estimation methods, (3) investigated a wide range of k-mer sizes, (4) applied a single, scalable ANI framework to diverse fungal phyla and oomycete taxa, large datasets, and understudied groups (e.g., oomycetes and non-Dikarya fungi), and (5) considered classification at the species rank as well as at the intraspecies and species-complex level. Our results show that sourmash-derived ANI provides accurate predictions of taxonomic assignments and phylogenetic relationships within fungal species, achieving performance comparable to conventional phylogenomic methods while offering substantially greater computational efficiency.

## Results

### Sourmash is more computationally efficient than conventional whole genome approaches

Sourmash was orders of magnitude more computationally efficient than a conventional whole genome ANI estimation method, ANIb, and a common orthologous gene phylogenomic approach in going from genome assemblies to a tree. For example, for 44 assemblies in the Pleosporaceae, it took 7 seconds to create genome sketches, compute comprehensive pairwise comparisons, and produce a sourmash ANI tree, while the ANIb approach was 2,720 times slower at 5.3 hours (Figure 1). Producing a maximum likelihood Benchmarking Universal Single Copy Orthologues (BUSCO) tree from 544 single copy orthologs took 38 hours and was thus 19,565 times slower than sourmash. Peak memory usage was lowest for sourmash at 120.8 Mb, compared to 534.4 Mb for ANIb and 41,830.2 Mb for BUSCO (Figure 1). Furthermore, it took 23 Mb of memory to store all 44 genomic signatures, while storing all 44 assemblies required nearly two orders of magnitude more memory at 1.4 Gb. In short, compared to ANIb and the BUSCO maximum likelihood approach, sourmash requires less memory storage, working primarily with signatures, and much less RAM while being orders of magnitude faster.

**Figure 1.**
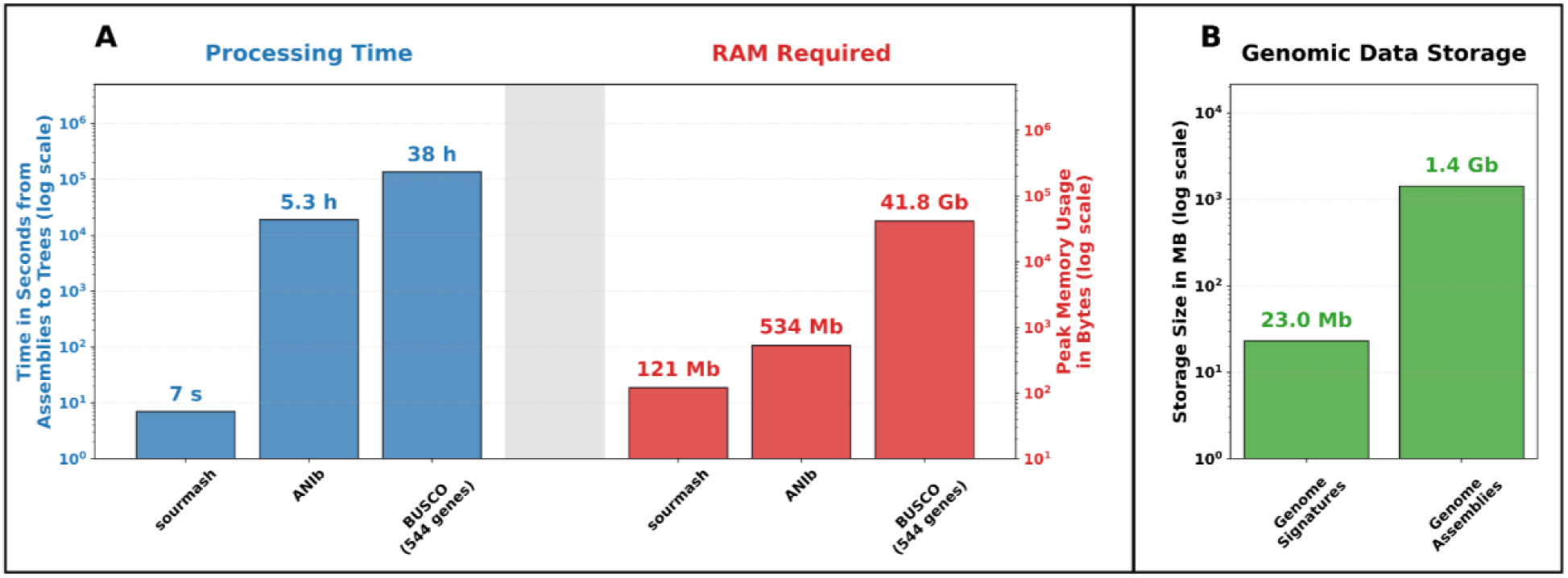
**Processing time, peak RAM** and data storage requirements for processing 44 genomes in the Pleosporaceae from assemblies to a tree via sourmash ANI, ANIb, and concatenated BUSCO gene methods, respectively. (A) The processing time and peak RAM required to compute a tree by sourmash ANI, ANIb, and using a maximum likelihood approach with a 544 BUSCO gene supermatrix. (B) Memory required to store the 44 genome assemblies compared to storing 44 sourmash genome signatures using a k-mer size of 21 and scaled value of 1000.

When comparing sourmash and ANIb results with each other, we found them to yield highly correlated ANI estimations across species, genus, and family ranks. When evaluated across a large dataset of fungal and oomycete assemblies (5,739 unique assemblies across 2,590 NCBI-defined species), both methods demonstrated strong agreement at lower taxonomic ranks (Supplemental Figure 1 and Supplemental Table 2). However, both methods lost accuracy for comparisons at the order rank and above as most ANI methods are not trustworthy below 75% similarity [36] and are not recommended for use below the 75-80% ANI range [35]. The speed of sourmash permitted more computations than ANIb and was independently assessed on even more genomes for a total of 16,228 unique assemblies (Supplemental Figure 2). Additional details on the ANIb benchmarking and implementation specifics are provided in the Supplementary Methods, Results, and Discussion.

### Sourmash ANI Accurately Recovers Fungal Taxonomy Below the Family Rank

After finding that sourmash and ANIb results were highly correlated, but sourmash was computationally much more efficient, we focused on sourmash-derived ANI for the rest of our study. First, to establish which sourmash parameters best recover currently understood taxonomy across the fungal and oomycete trees of life and at different taxonomic ranks, we evaluated parameter performance across 13 diverse families spanning four phyla. We focused on two oomycete families (Peronosporaceae and Pythiaceae), three families in the Mucoromycota (Mortierellaceae, Mucoraceae, and Lichtheimiaceae), four families in the Ascomycota (Cordycipitaceae, Hypocreaceae, Nectriaceae, and Pleosporaceae), and four families in the Basidiomycota (Boletaceae, Polyporaceae, Trichosporonaceae, and Ustilaginaceae). Tanglegram comparisons revealed high topological concordance between k21, max containment sourmash ANI trees and core-genome BUSCO trees across family, genus, and species ranks (Figure 2). No isolates were misclassified at the species rank in sourmash trees when considering clades in the core genome trees as ground truth. A species rank legend for Figure 2 can be found in Supplemental Figure 3, and the number of common single-copy BUSCO genes utilized for producing reference trees are displayed in Supplemental Table 4.

**Figure 2.**
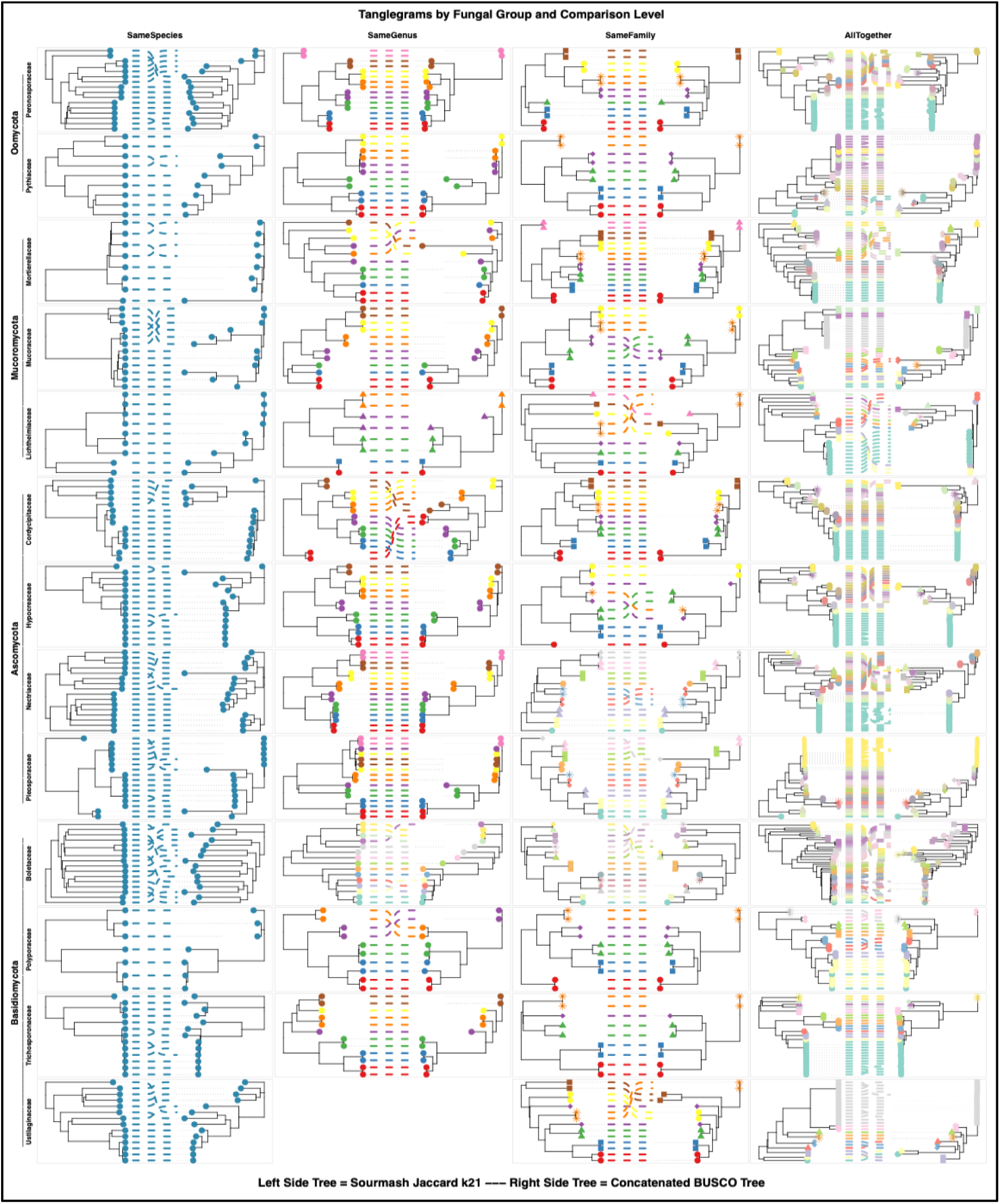
Concordance between sourmash ANI trees and concatenated single copy ortholog trees for oomycete and fungal families in various phyla. Tangelgram so sourmash-derived ANI trees (left side trees) vs. concatenated BUSCO trees (right side trees) at various taxonomic ranks (species, genus, and family). The sourmash trees use a k-mer size of 21 and max containment for whole genome similarity estimation.

Robinson-Foulds (RF) distance metric analysis demonstrated that k-mer size impacts concordance between sourmash and BUSCO trees differentially depending on the taxonomic rank being evaluated (Figure 3). We quantified weighted and unweighted RF distance across ranks of family, genus, and species to establish concordance between BUSCO-based trees and sourmash trees created using both Jaccard and max containment-inferred ANI for sourmash signatures computed using k-mer sizes of 16, 21, 31, 51, 61, and 71 (histograms of RF values in Supplemental Figures 4 and 5). Mean and standard deviation of RF distances per rank are shown in Figure 3, and k16 results were left out of the trendline and correlation metrics as they did not follow the same trend as the other k-mer sizes. At the family rank, topological discordance scaled positively with k-mer size, as weighted RF distances increased from k21 to k71 with strong correlation and steep slopes of 7.4×10^−3^ and 7.8×10^−3^ for Jaccard and max containment, respectively (Figure 3). Sensitivity to degradation of tree agreement for larger k-mers decreased at the genus rank with lower correlations and shallower slopes, and slope dropped lowest during same-species comparisons. Similar patterns are seen in the bottom half of Figure 3 with unweighted RF distances, where the slope is greater at the same family rank compared to same genus and species rank comparisons and with correlation between k-mer size and RF distance decreasing as taxonomic distance decreases. While Jaccard-inferred ANI achieved marginally lower mean RF distances than max containment ANI across in majority of evaluations (12/18 weighted and 11/18 unweighted mean distances), their standard deviations highly overlapped, indicating largely equivalent performance (Figure 3). RF distance analysis shows that k21 is the most robust k-mer size across all taxonomic ranks considered, while larger k-mer sizes like 51, 61, and 71 produce ANI trees less concordant with BUSCO trees at the family rank but increase in accuracy as taxonomic scope decreases from the family to the species rank.

**Figure 3.**
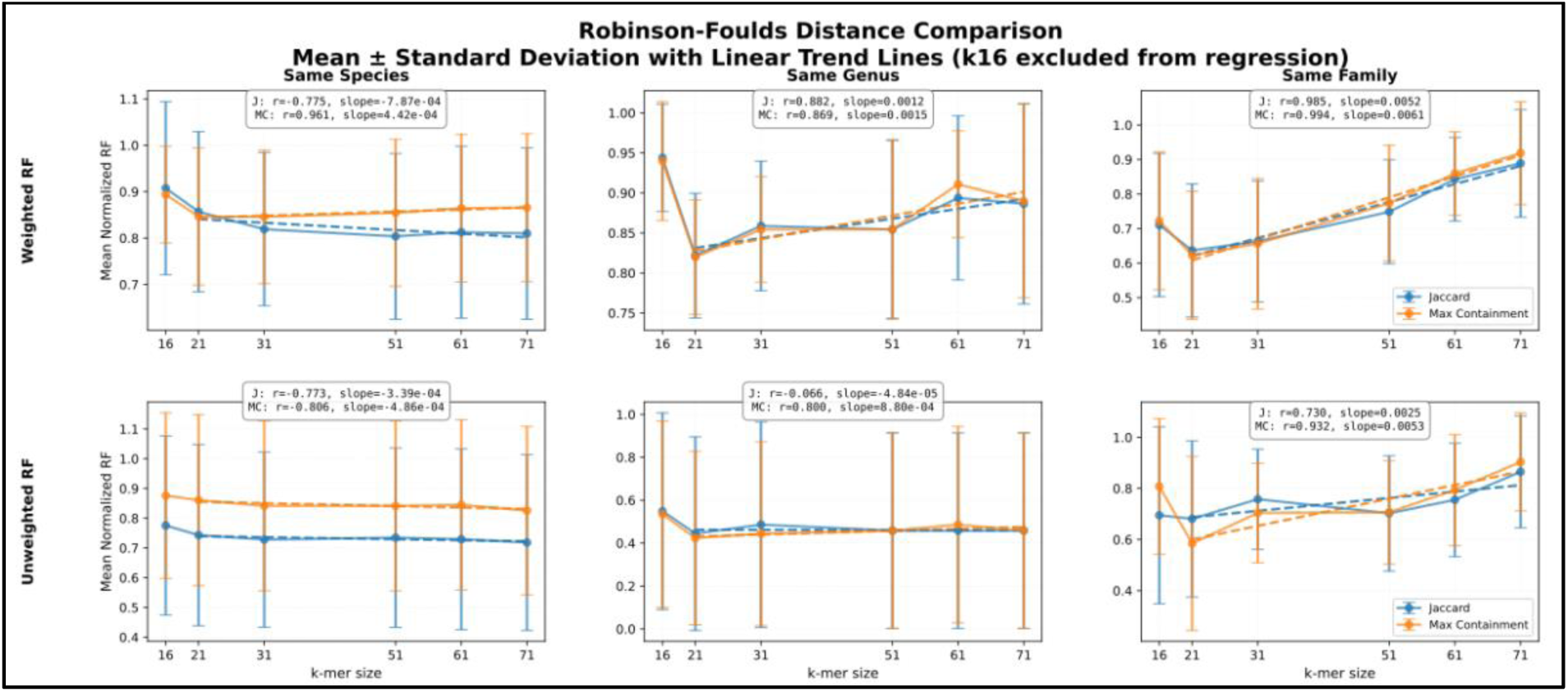
Tree distance metrics estimating agreement between sourmash ANI and concatenated single copy ortholog trees for fungal and oomycete taxa at varied taxonomic scope including isolates from the same species, same genus, and same family. Weighted Robinson-Foulds (top row) and unweighted Robinson-Foulds (bottom row) tree distance metrics analysis comparing core genome concatenated BUSCO phylogenies to sourmash ANI-based trees created at using various k-mer sizes (k = 16, 21, 31, 51, 61, and 71) and ANI estimation methods (Jaccard similarity and max containment). Data points and error bars represent the mean and standard deviation across all fungal and oomycete families for a specified k-mer size and ANI estimation method as compared to the BUSCO tree. The dotted lines represent the linear regression line of best fit for k-mer sizes ranging from 21 to 71 (k16 is left out of this computation) and the Pearson’s r and slope values are displayed for each plot.

### Whole genome similarity resolves species for varied taxa of fungi and oomycetes

Whole genome similarity, as determined using sourmash, was next assessed for its ability to successfully classify larger clades of isolates compared to conventional taxonomic and phylogenomic methods in a variety of fungal and oomycete groups with varied taxonomic scopes ranging from the order to genus ranks. Sourmash ANI-estimation methods achieved perfect species-rank classification accuracy across a phylogenetically diverse suite of fungal and oomycete taxa evaluated in datasets at scopes ranging from the genus to order levels. Based on the prior analysis and given the multi-genus scope of the trees considered, a k-mer size of 21 was utilized for the following analysis. Benchmarking both Jaccard and max containment similarity estimation methods revealed zero species-rank misclassifications compared to reference BUSCO trees across all groups evaluated, with max-containment ANI topologies illustrated in Figure 4. Two genera in the fungal order Onygenales populate the tanglegram in panel A with a sourmash and a 329-gene BUSCO tree and all species in both trees form monophyletic clades. Figure 4B displays a few families in the oomycete order Saprolegniales with a sourmash tree compared to a 31-gene BUSCO maximum likelihood tree. All species formed the same species rank clades in the sourmash tree as they did in the BUSCO tree, but there was a difference in branching for isolates in different families. The other oomycetes in Figure 4, *Phytophthora* clades 8 and 1 in panels C and D respectively, show similarly high concordance between whole genome similarity trees and core genome trees (produced with 26 and 22 single copy genes for clades 1 and 8 respectively). Figure 4E displays a tanglegram with a 76-gene BUSCO tree for members of the fungal family Pucciniaceae, a group high in transposable element content, and species rank clade membership is the same between the two trees. The same Pucciniaceae tanglegram can be seen in Supplementary Figure 6 with total transposable element percentages estimated for 45 of 52 assemblies by EarlGrey noted ranging from 22.0% to 76.1%. Panel F displays a Mortierellaceae family tanglegram in which certain NCBI-defined species appear to not form monophyletic clades; however, each clade contains the same isolates in the sourmash as the BUSCO tree produced with 132 genes. The high concordance between sourmash ANI trees and BUSCO orthologous gene trees further confirms the accuracy of species rank classification achieved using sourmash.

**Figure 4.**
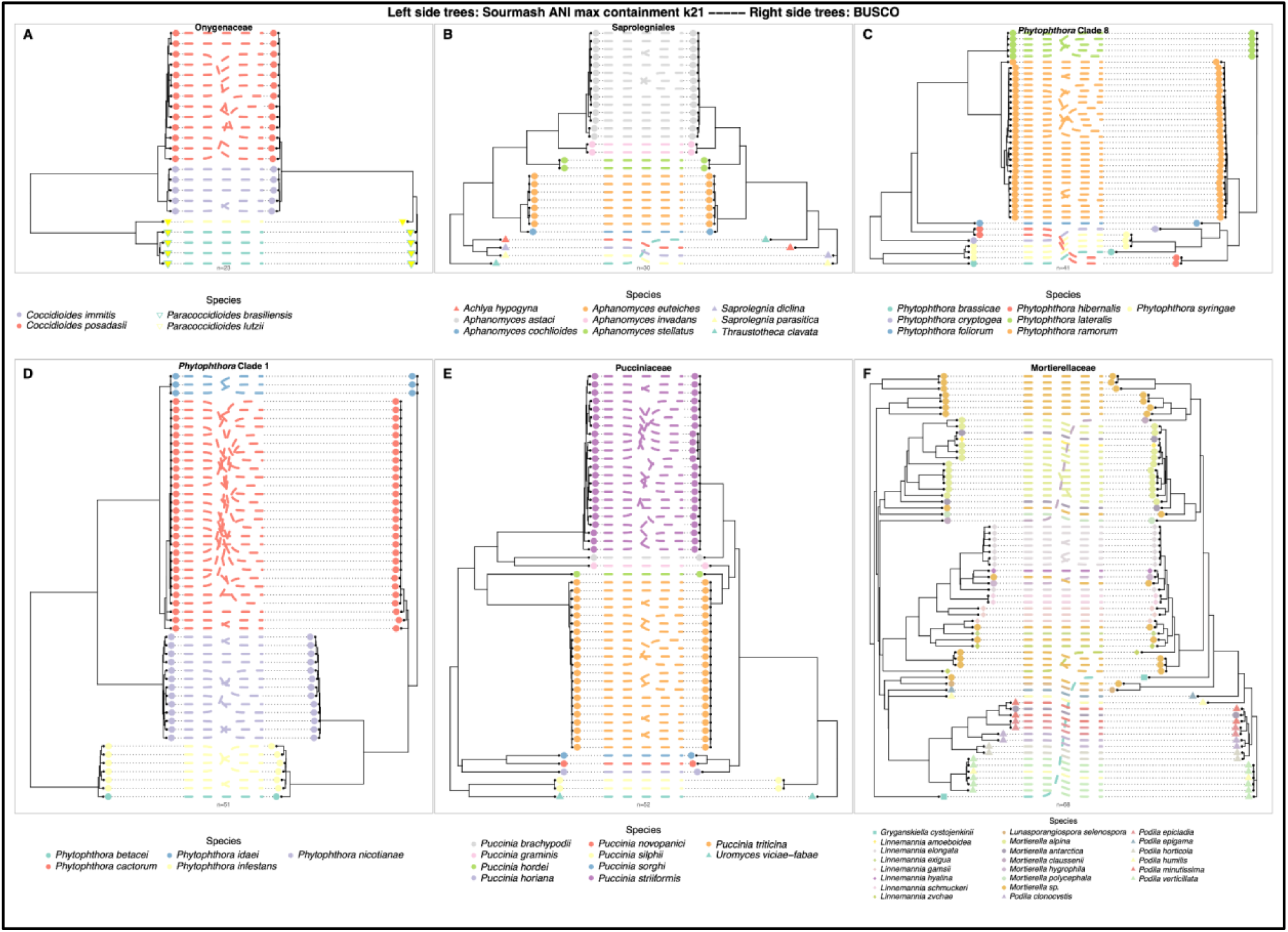
Species rank clade concordance between sourmash ANI trees (left side trees) and concatenated BUSCO gene maximum likelihood trees (right side trees) for isolates from the following taxa: (A) Onygenaceae, (B) Saprolegniales, (C) *Phytophthora* clade 8, (D) *Phytophthora* clade 1, (E) Pucciniaceae, and (F) Mortierellaceae.

### Whole genome similarity for species complex classification

Sourmash successfully resolved species and lineages within fungal species complexes, matching reference orthologous gene tree clades and resolving NCBI curation conflicts. *Rhizopus* and *Cryptococcus* ANI matrices were calculated using a k-mer size of k21 given the multi-species nature of these trees; while, for the *Boletus edulis* tree, which contains primarily only members within the complex, a variety of k-mer sizes were tested ranging from k16 to k71. Jaccard and max containment similarity estimation methods were used to consider which achieved the fewest misclassifications at the species rank for *Rhizopus* and *Cryptococcus* and the lineage level for *Boletus edulis*.

The *Rhizopus microsporus* and *R. arrhizus* complexes, along with *R. stolonifer* and members of *Mucor*, formed the same species clades in a sourmash tree as determined by a 602-gene BUSCO tree (Figure 5A). *R. microsporus* variants as defined in NCBI do not all land in monophyletic clades; however, all variants share the same clade membership between the two trees. Similarly, the *R. arrhizus* complex, including *R. arrhizus* and *R. delemar,* formed clades all containing the same isolates between the two trees while some NCBI species designations did not for monophyletic clades yet agree with isolate placement between the trees. The *Cryptococcus* tanglegram in Figure 5B shows high concordance between a 396-gene BUSCO tree and sourmash tree. All assemblies form the same monophyletic clades between the two trees despite isolates with conflicting NCBI species designations. Neither *Rhizopus* nor *Cryptococcus* trees had any species rank misclassifications.

**Figure 5.**
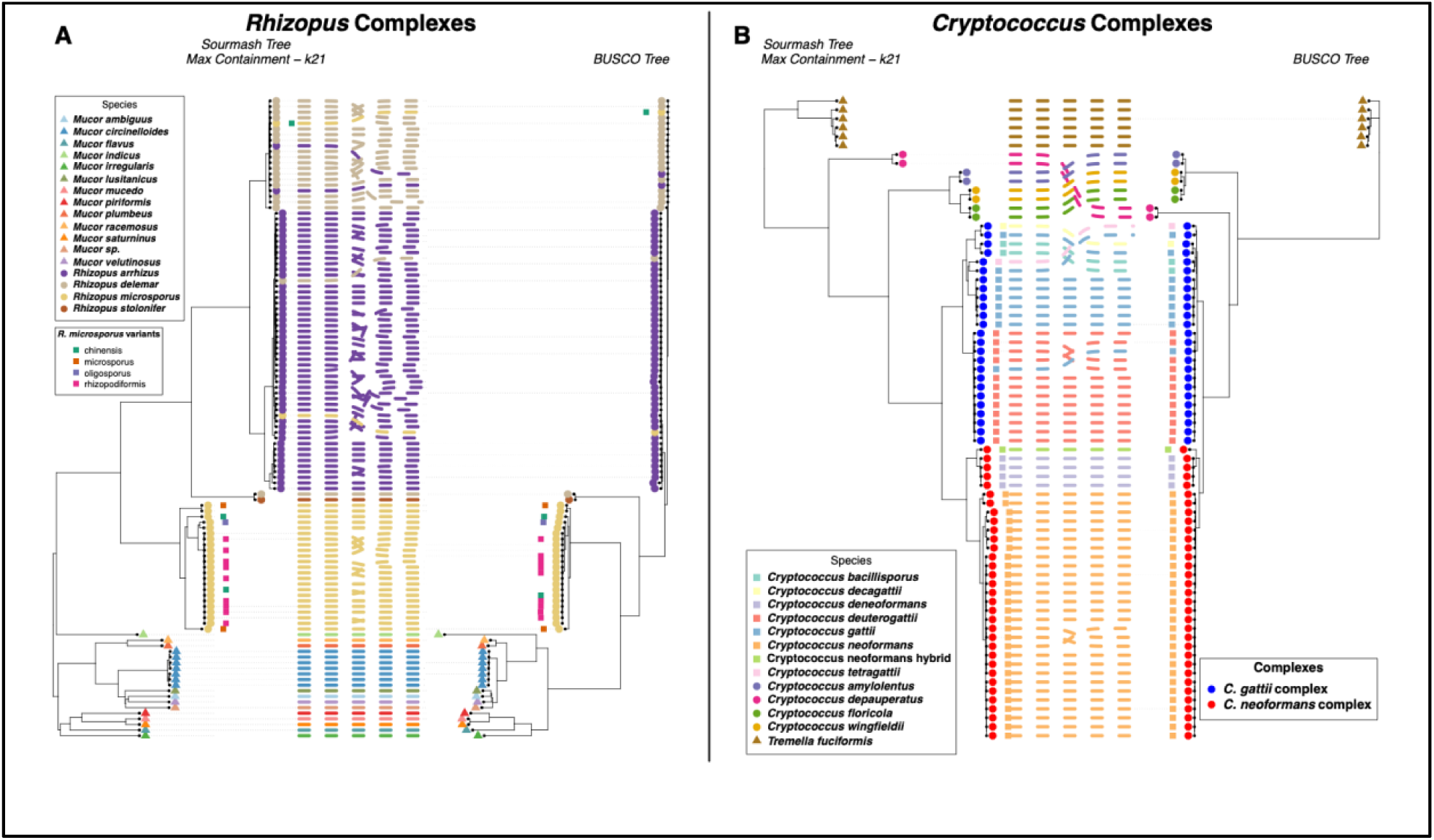
Concordance between sourmash ANI trees (left side trees) and concatenated BUSCO gene maximum likelihood trees (right side trees) displaying species and species complex level clade agreement between methods for isolates from the following taxa: (A) Members of the Mucorales including the *Rhizopus arrhizus* and *R. microsporus* complexes, and (B) Members of the Tremellales including the *Cryptococcus neoformans* and *C. gattii* complexes.

Within the *Boletus edulis* species complex, lineage classification was optimized by utilizing max containment and larger k-mer sizes (k≥51), which minimized subspecific misclassifications as defined in a coalescent orthologous gene tree reference. Lineage level misclassifications for *B. edulis* were greater for Jaccard-based ANI (3 to 11 misclassifications) than max containment-based ANI (1 to 3 misclassifications) and were generally greater for smaller k-mer sizes like k21 and k31 (2 to 9 misclassifications) compared to larger k-mer sizes like k51 through k71 (from 1 to 4 misclassifications) (Supplementary Table 5). High concordance between the reference tree and ANI tree is displayed in the Figure 6 tanglegram with the sourmash tree created using max containment and a k-mer size of 51 (which achieved the fewest misclassifications along with max containment k16, k61, and k71). In this case, of the 252 *B. edulis* isolates in the trees, a single isolate in the EC lineage in the sourmash tree did not cluster with all other isolates of its lineage to form a monophyletic clade. In conclusion, sourmash ANI was highly accurate in classifying species and members of species complexes.

**Figure 6.**
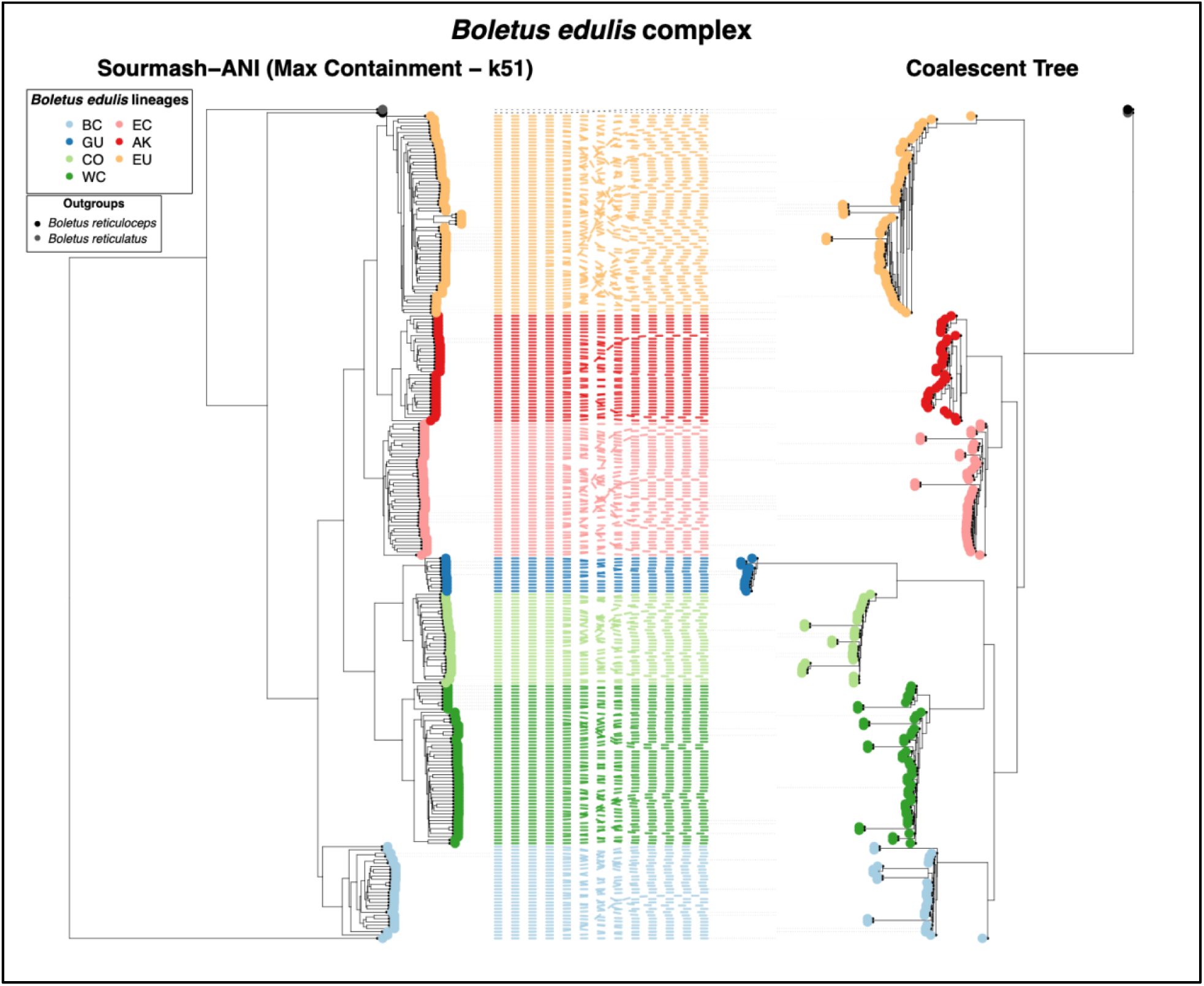
Concordance between a sourmash ANI tree (left side trees) and summary coalescent gene tree (right side trees) displaying clade agreement between lineages in the *Boletus edulis* complex with *Boletus reticulatus* and *B. reticuloceps* as outgroups.

### Whole genome similarity for intraspecific variant classification

Intraspecific variant classification via sourmash ANI was highly accurate with respect to SNP-based reference topologies, especially when utilizing larger k-mer sizes. The accuracy of sourmash to classify subspecies designations at various k-mer sizes and ANI estimation methods was assessed in four high-consequence fungal and oomycete pathogens including the ascomycete species *Aspergillus fumigatus* and *Magnaporthe oryzae*, the chytrid *Batrachochytrium dendrobatidis*, and the oomycete *Phytophthora ramorum*. Quantitative evaluation of weighted and unweighted RF distances showed that large k-mer sizes (k≥51) consistently generated subspecific trees with the highest concordance to reference tree topologies (Figure 7E, Supplementary Figure 7). Tanglegrams generated using the top-performing unweighted RF parameters (k=71 for *A. fumigatus*, *M. oryzae*, and *P. ramorum*; k=61 for *B. dendrobatidis*) revealed strong clade agreement between SNP-based and sourmash trees (Figure 7).

**Figure 7.**
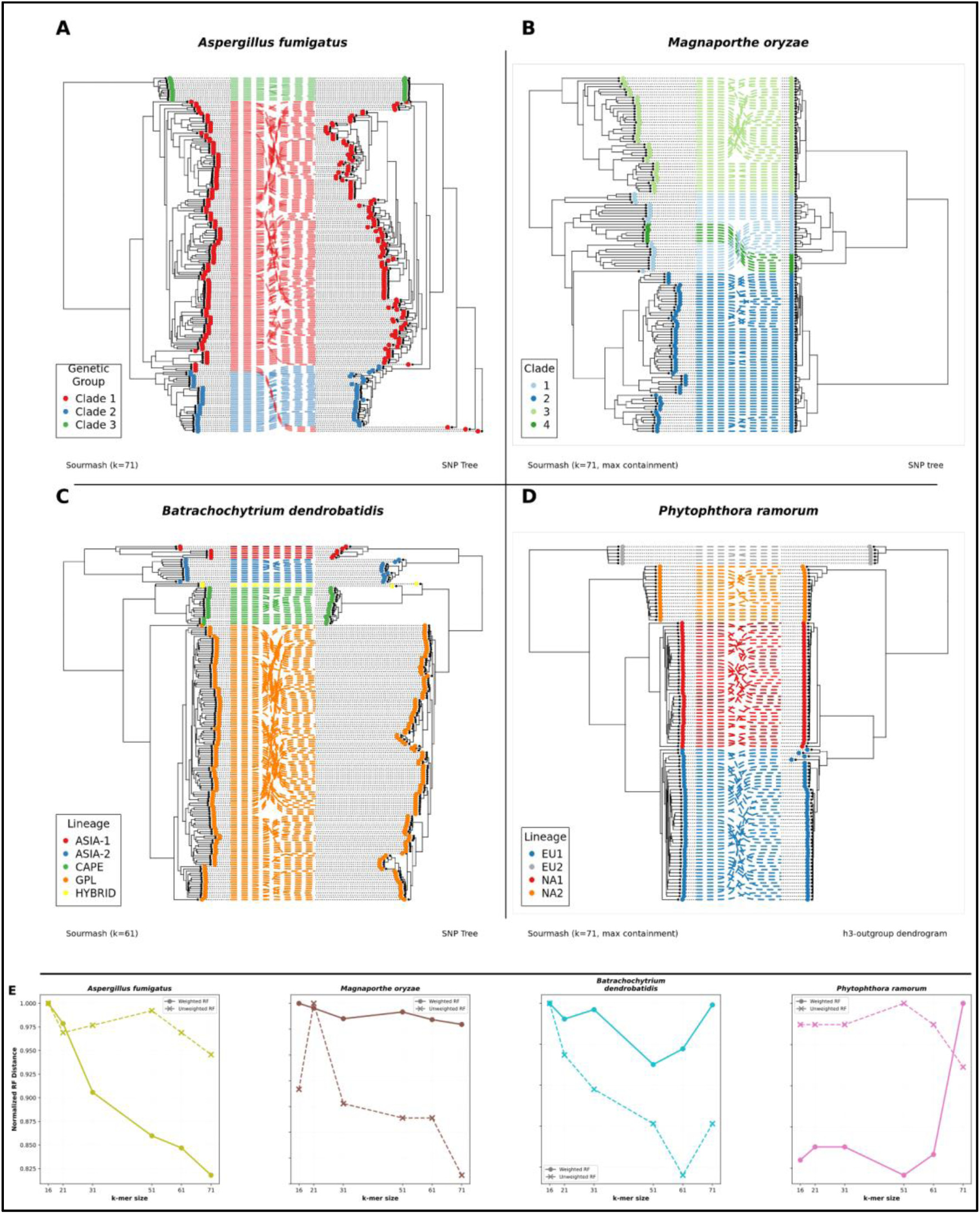
Concordance between sourmash ANI trees (left side trees) and SNP-based dendrograms (right side trees) displaying subspecies level clade agreement between methods for isolates from the following fungal and oomycete species: (A) *Aspergillus fumigatus*, (B) *Magnaporthe oryzae*, (C) *Batrachochytrium dendrobatidis*, and (D) *Phytophthora ramorum*. (E) Normalized RF distances computed between SNP-based trees and max containment sourmash ANI trees at various k-mer sizes from k16 to k71.

All *A. fumigatus* intraspecific genetic groups form monophyletic clades in the sourmash max containment ANI tree despite clade 1 not being monophyletic in the SNP tree (Figure 7A). In *Magnaporthe oryzae*, sourmash max containment ANI accurately recovered three of the four infraspecific lineages but split clade 1 into four distinct sub-clusters compared to the reference f3-outgroup dendrogram (Figure 7B). *B. dendrobatidis* lineages all form clades with the same intralineage clade membership for both sourmash and the reference SNP trees (Figure 7C). For *Phytophthora ramorum*, sourmash max containment ANI achieved perfect subspecific classification, resolving all four well-characterized lineages into distinct monophyletic clades that mirrored the clades resolved in the reference biallelic SNP tree (Figure 7D). Based on this analysis, sourmash ANI is sufficient for subspecies rank designations in the taxa assessed and larger k-mer sizes may be best suited for intraspecific classification using sourmash.

Max containment-inferred ANI achieved higher lineage-level classification accuracy by clade membership than Jaccard, yielding fewer subspecies lineage misclassifications across all k-mer sizes for *P. ramorum* and all but k21 for *A. fumigatus* (Supplementary Table 5). Several misclassified isolates in these taxa exhibited anomalies in assembly size compared to the rest of the dataset. The single isolate misclassified by Jaccard-based ANI in the *P. ramorum* dataset had a genome size of 88 Mbps while the remaining assemblies ranged from 53-70 Mbps. Similarly, two assemblies misclassified by Jaccard-based ANI in the *A. fumigatus* dataset had genome sizes of 38 and 61 Mbps while the remaining assemblies are in the range of 25-30 Mbps. Max containment-inferred ANI also had fewer misclassifications for *B. dendrobatidis* at smaller k-mer sizes of k16-k31; while both methods tied for 0 misclassifications at k-mers ranging from k51-k71 (Supplementary Table 5). Jaccard-based ANI had fewer misclassifications than max containment-based ANI for *M. oryzae* for all k-mer sizes except k51 where the methods tied. Overall, max containment-inferred ANI achieved higher lineage level classification accuracy by clade membership.

## Discussion

Whole genomes provide a wealth of information for identification of fungi and oomycetes, a critical task in biological, agricultural, and medical sectors; however, efficient computational methods are required to leverage whole genomes for taxonomic classification at scale. This study approaches this problem by systematically evaluating the accuracy and efficiency of a rapid, FracMinHash sketch-based genome similarity method for fungal and oomycetous taxonomic categorization. Compared to modern gold standard concepts in fungal taxonomy such as phylogenomic-based circumscription requiring hundreds of orthologs or massive SNP datasets, sourmash ANI recovers genus, species, and subspecies-level groupings with high concordance but with greater speed and a fraction of the computational resources.

The current study, to our knowledge, represents the largest collection of fungal assemblies assessed by ANI to date. Prior whole genome similarity investigations in fungi overwhelmingly use alignment [9–23] or mapping-based approaches [24, 25, 37–41], and most are limited to a single taxon or taxonomic group. We expand ANI-based fungal taxonomy to alignment-free, sketching methods and extend ANI analysis beyond the Dikarya to relatively understudied groups like members of Chytridiomycota, Mucoromycota, and Oomycota. The few multi-phylum studies of fungal whole genome similarity focus on the similarity boundary between species as defined in databases [22, 24, 39, 42] or assembly level database querying [32]; whereas we benchmark clade-level taxonomic placement against reference phylogenomic trees. By extending analysis to species complexes and subspecific ranks we show a single whole genome similarity framework can remain informative across evolutionary scales from the genus rank through infraspecific lineages. Furthermore, we compared sketch-based ANI to conventional BLAST-based ANI and assessed biological and technical considerations of k-mer-based distance estimations in fungi, such as assembly quality and sourmash ANI estimation parameters.

One striking result of this study was the degree to which species- and lineage-level clustering inferred from FracMinHash sketch comparisons matched relationships as defined by ortholog and SNP-based phylogenomic approaches despite the fundamentally different information content used by the methods. BUSCO and SNP approaches focus on specific genomic regions, whereas sourmash-based ANI summarizes genome-wide sequence similarity. Agreement between ANI trees and reference phylogenies was observed across Ascomycota, Basidiomycota, Mucoromycota, Chytridiomycota, and Oomycota.

In working to develop best practices for community standards, we assessed the accuracy of sourmash ANI at classifying broad fungal diversity at various taxonomic ranks using a range of k-mer sizes from k16-k71. We found that overall, k21 is sufficient for accurately classifying fungal and oomycetous genera and species and is the most robust k-mer size across taxonomic ranks considered in this study. There was a trend of improved performance for larger k-mer sizes as taxonomic distances decreased, and through systematic evaluation of intraspecific variation in this framework, k51-k71 were most accurate in the taxa assessed. Jaccard similarity and max containment ANI estimation approaches generally achieved similar mean RF distances between sourmash and reference trees in the datasets considered, but max containment achieved fewer misclassifications of isolates in datasets with a wide range of genome sizes and low BUSCO completeness. This result reflects the resilience of containment metrics to assembly size and is consistent with a prior study noting the robustness of max containment to disparate assembly sizes [43]. Consequently, our findings suggest that max containment-inferred ANI is a robust choice when working with assemblies of fluctuating quality and coverage, or in biological systems where the goal is to minimize the impact of large insertion events which may penalize accuracy of Jaccard-based classification. Additionally, every isolate which appeared within a clade of isolates bearing another species name (i.e., isolates which likely have the incorrect species currently assigned in NCBI) appeared in the same clades in reference trees as in sourmash trees, therefore sourmash ANI can be used as a method for flagging potentially mis-identified isolates in large databases where conventional methods would be cumbersome.

Despite reducing genomes to compact sketches representing a small fraction of their total content, sourmash consistently recovered accurate taxonomic groupings across fungi and oomycetes. A scaled value of 1,000 was used for all analyses, meaning 1 in 1,000 unique k-mers was kept per genomic assembly to create each signature. It’s possible that a larger scaled value, which would reduce the signature size, could still achieve high accuracy while compressing data further. A scaled value of 1,000 matches what sourmash developers recommend for bacterial and archaeal sketches and metagenomic analysis [44]. As fungal assemblies are generally larger, a greater number of k-mers are retained per sketch compared to prokaryotes at a selected scaled value. Sourmash analyses require the scaled value of a query be the same or smaller than a target sketch to maintain accuracy; thus, using a scaled of 1,000 for fungal genomes ensures the sketches are compatible with prokaryotic sketches for applications in metagenomic profiling. The compression of assemblies leading to orders of magnitude reduction in memory required for storage makes sourmash an attractive option for large scale databases compared to storing full genomic assemblies.

Whole genome similarity-based taxonomy using sourmash was orders of magnitude more efficient than a BUSCO gene or an ANIb approach, making it an attractive option for taxonomic classification as whole genomes continually become cheaper and easier to produce causing genomic database sizes to increase exponentially, especially considering sourmash has further functionality for database searches such as indexed sequence bloom trees for working with large signature databases [45] which are not feasible for conventional phylogenomic or ANIb methods. As sourmash ANI can classify fungi in seconds, a sourmash taxonomic classification scheme would be fast enough to open new avenues for large-scale genome collection and analysis in research and industry with reduced computational and economic costs. This speed accompanied by a relatively lower degree of bioinformatic expertise for ANI classification compared to a BUSCO or SNP-based approach can help lower the bar of experience required for genomic classification of fungi and may benefit applications like sequence analysis for non-specialists, agriculture, hospitals, or community science. Another benefit of a whole genome ANI approach includes the ability to rapidly add an isolate to an existing tree. For a single copy ortholog phylogeny, the whole tree building workflow must be repeated to add a single isolate; whereas a whole genome similarity approach only requires comparing the newly considered genome assembly to all prior assemblies before computing a new tree.

All taxa that we considered were resolved to the species rank using this whole genome similarity approach. However, sourmash ANI trees exhibited some branching differences above the genus rank compared to BUSCO gene maximum likelihood trees while achieving high-quality clustering of species, reinforcing that ANI is beneficial as genus, species, and subspecies classification tool rather than a tool for deep phylogenetic inference. In assessing species complexes, *Rhizopus microsporus* variants did not all form monophyletic clades, and this result agrees with a recently published SNP-tree which found variants as currently defined to be paraphyletic [39]. The most notable subspecies level discrepancy between sourmash ANI and reference trees observed was in *Magnaporthe oryzae* where clade 1, the only sexually recombining group among 3 clonal lineages [35], did not form a monophyletic clade as in the reference f3-outgroup dendrogram and formed a sister clade to clonal clade 4. We analyzed the *M. oryzae* dataset further to assess how other methods cluster isolates, and both a BUSCO and a FastANI approach produced trees where clade 1 was not a monophyletic clade (Supplemental Figure 8 and 9). Clade 4 and a portion of clade 1 being sister clades in the sourmash ANI tree, as well as other relationships in the tree, could be due to the presence of accessory chromosomes which are known to be present in many of the isolates considered [46], and it could be expected that a whole genome similarity approach may be sensitive to accessory genome content while a SNP-based method like an f3-outgroup dendrogram may be less sensitive to accessory chromosomes. While containment metrics limit unshared sequence content from inflating taxonomic distance across the entire dataset, the unique k-mer content contributed by accessory content may still contribute to the overall ANI computation and more so for Jaccard methods. Sensitivity to accessory content could prove valuable in fungal genomics where a core genome focused approach may not consider accessory, horizontally acquired, or rapidly evolving genomic content which may be relevant to phenotypic traits like pathogenicity.

The only species rank discrepancy between reference and sourmash ANI trees was the single isolate of *Boletus edulis* in the BC lineage which landed on an isolated long branch in the ANI tree, outside of the *B. edulis* clade further from *B. edulis* than *B. reticulatus* or *B. reticuloceps*. The isolate in question had a very low BUSCO completeness of 3.1% which is likely the reason for its suboptimal placement in the ANI tree, despite other isolates with BUSCO completeness of 9.9%, 12.1%, and 13 others in the range of 27.3-49.7% all landing in their appropriate clades in the ANI tree [47]. This result, along with the *Aspergillus fumigatus* and *Phytophthora ramorum* lineage classifications, highlights the robustness of the sourmash max containment ANI approach to a broad range of genome completeness and lengths. Moving forward, it will be essential to consider how genome quality and biological attributes such as repetitive content, hybridization, ploidy, heterokaryosis, and accessory chromosomes influence accuracy and are incorporated into standard development for whole genome-based classification methods.

Fungal taxonomy is rapidly changing, and this instability has implications for medical mycology, plant pathology, and food safety, in which rapid and precise fungal identifications guide decisions on response efforts with human health repercussions. Because functional traits are associated with groups of related fungi, better understanding their relationships using tools like these will likely offer novel insights into products and processes that can be developed in applied settings. Lastly, for fundamental biological and evolutionary research, the clarity of results and the associated conclusions may be skewed if researchers are comparing fungal genomes of isolates without understanding the genomic divergences between them. We argue that standardized ANI methods provide a high throughput means of taxonomic classification that complements, rather than replaces, traditional data such as sexual compatibility, ecological, geographical, and morphological data.

## Conclusion

Whole genome similarity-based classification using sourmash is orders of magnitude more computationally efficient than ANIb and phylogenomic approaches while achieving comparable accuracy across species and subspecies ranks. A k-mer size of 21 was most robust for species rank classification while larger k-mer sizes (51–71) were optimal for intraspecific resolution, and max containment-inferred ANI was more robust to genome size variation and completeness than Jaccard. These results demonstrate that scalable, alignment-free approaches can enable genome-based taxonomy across the rapidly expanding diversity of fungal and oomycete genomes. Overall, this work establishes whole genome similarity determined using FracMinHash sketch-based methods as rapid and effective for taxonomy for a broad range of fungi and oomycetes and supports the development of large-scale genomic reference databases and standardized, rapid identification pipelines for research, clinical, and agricultural applications.

## Methods

### Data sources and assembly methods

Genome assemblies and associated metadata were primarily obtained from the NCBI Assembly database using the NCBI Datasets tool (v. 18.3.1). Freely available large language models including Claude, Gemini, Grok, and DeepSeek were utilized to dynamically help write and edit the code utilized throughout this manuscript (January 2025–May 2026). A comprehensive metadata table including species names and accession numbers for all NCBI fungal and oomycete assemblies was downloaded on April 14, 2025 and used for multiple downstream analyses. Metadata was filtered to remove all species with genus designations in brackets, all species crosses denoted by an ‘x’, ‘aff’ or ‘cf’ in the species name, and all GCF accessions. Taxonomic rank information was assigned using the NCBITaxa class implemented in ete3 (v. 3.1.3) in Python (v. 3.8.15). Metadata processing and filtering were performed using pandas (v. 2.0.3). For additional isolates beyond publicly available assemblies, raw sequencing reads were obtained from the NCBI Sequence Read Archive (SRA) using the fasterq-dump function of the SRA Toolkit (v. 3.1.1). Unless stated otherwise, reads were filtered using fastp (v. 0.24.0) applying the flags ‘--qualified_quality_phred 20’ and ‘--length_required 50’ for all reads and additionally the ‘--detect-adapter-for_pe’ flag for paired-end reads followed by SPAdes (v. 4.1.0) for genome assembly with the ‘--cov-cutoff auto’ flag. The Entrez submodule in the python Bio (v. 1.83) package was used to determine the nature of the read files (sequencing tech, single vs. paired end, etc.). Previously published genome assemblies and tree data were incorporated where necessary and are referenced accordingly. Descriptions of dataset composition are provided within each analysis-specific methods and results section.

Tree building and assessment methods

Sourmash (v. 4.8.4) was used to create genome sketches and perform pairwise ANI computations using both Jaccard similarity and max containment, with a scaled value of 1,000 for all signatures meaning that on average 1 k-mer was retained per 1,000 unique k-mers per assembly. To produce sourmash trees, ANI matrices of all pairwise comparisons per dataset were converted to distance matrices by computing 1 minus the ANI matrices. Distance matrices were used to compute unweighted pair group method with arithmetic mean (UPGMA) trees using the Bio package in Python. Tanglegrams were produced to visually compare sourmash and reference trees using the ape package (v. 5.8.1) to read tree data and cophylo package (v. 2.4.4) to create the tanglegrams in R (v. 4.3.3).

Orthologous gene phylogenies were produced by identifying single copy orthologs using the BUSCO (v. 5.8.3) software to annotate single copy orthologs, and the BUSCO databases used in each analysis are noted in their corresponding sections. For each dataset all common single copy BUSCO genes were retained. Each retained BUSCO gene family was aligned using MAFFT (v. 7.526) with the “--auto” flag and “--maxiterate” set at 1000, all genes per genome assembly were concatenated, trimmed using trimAl (v. 1.5.rev0) with the ‘-strictplus’ flag, and maximum likelihood trees were produced using iqtree2 (v. 2.4.0) using the ‘-MFP+MERGE’ flag, “-bb’ set to 1000, and “- alrt’ set to 1000. All BUSCO trees in this manuscript were created using this method unless otherwise stated. Phylogenetic trees were further processed using the Bio package in Python with the phylo submodule.

Sourmash ANI trees were evaluated by comparing clade membership between ANI and BUSCO or SNP-based phylogenies and computing weighted and unweighted RF distances between the sourmash and reference trees - computed using the dendropy (v. 5.0.1) package in Python. RF distances were computed using the ‘dendropy.calculate.treecompare’ submodule, and all tree branch lengths were normalized by dividing each branch length by the total branch length of each tree prior to computing RF distances. Accuracy of sourmash trees was further assessed by considering the number of misclassified assemblies, with a misclassification defined as an assembly in the sourmash tree that appears outside of the largest monophyletic clade for a specific taxon as defined in the reference BUSCO or SNP-based tree.

### Sourmash k-mer size and ANI method analysis

Sourmash sketches and pairwise ANI were determined using k-mer sizes of 16, 21, 31, 51, 61, and 71bp. To evaluate how parameter choice affects dendrograms produced by sourmash ANI matrices, weighted and unweighted RF distances were quantified between sourmash ANI based trees and BUSCO-based maximum likelihood trees produced with isolates encompassing varied taxonomic ranks including family, genus, and species. Genome assemblies were selected to cover a broad range of fungal and oomycetous taxa while considering only families with enough assemblies in NCBI to have at least 8 members of one species and at least 4 pairs of assemblies from that genus and that family. Minor exceptions were made to increase the taxonomic breadth of this dataset.

A list of assemblies used in this analysis and all sourmash and benchmarking universal single copy orthologs (BUSCO) trees for this analysis can be found in the project’s GitHub and Zenodo repositories: https://github.com/Uehling-Lab/SourmashForFungalTaxonomy/tree/main/SupplementalData_Accessions_Trees_And_BUSCOdatabases/FamilyGenusSpecies_SourmashANIParameterAnalysis (DOI: 10.5281/zenodo.21707559), and the BUSCO database used for each family and number of common BUSCOs used to prepare orthologous gene trees are listed in Supplementary Table 4. Assemblies were downloaded from the NCBI assembly database except for the following: No genus rank tree was produced for the Ustilaginaceae due to the limited number of genome assemblies and read files available in NCBI. At the species rank of the Boletaceae, genome assemblies of *Boletus edulis* were acquired from Tremble et al. [47]. At the genus rank for Lichtheimiaceae, 9 genomes of *Lichtheimia* species including *L. blakesleaana*, *L. ornata*, and *L. hyalospora* were assembled from paired-end Illumina reads acquired from the NCBI SRA database. To detect potential contamination of genomes assembled from SRAs, initial assemblies were indexed, mapped, and sorted using bwa (v. 0.7.19-r1273) and samtools (v. 1.23) and binned using metabat2 (v. 2:2.18) with default parameters. Resulting bins were identified taxonomically with kraken2 (v. 2.1.6) using kraken’s ‘core_nt’ database downloaded on September 16, 2025, and bins were recombined into assemblies based on the top family rank hit. BUSCO completeness was then determined using the fungi_odb10 database, and the highest completeness genome pairs per species were retained.

For each species, genus, and family rank dataset, sourmash sketches were generated, followed by pairwise ANI estimation using Jaccard and max containment, and weighted and unweighted RF distances were computed between sourmash trees and BUSCO trees. RF distances were normalized by dividing RF distances by the highest RF distance achieved for each family and taxonomic rank combination. To determine the best sourmash parameters, we compared means of normalized RF distances for each parameter set at each taxonomic rank.

### Sourmash and ANIb computational efficiency comparison

To assess the speed and memory requirements of whole genome taxonomic approaches, sourmash, ANIb, and BUSCO were run for all 44 genome assemblies in the Pleosporaceae used in the preceding analysis (sourmash k-mer size and ANI method analysis). Sourmash signatures were computed with k21 and max containment was used as the ANI estimation approach. The software pyANI (v. 0.2.12) was used for ANIb computations, and ANIb matrices were used to produce UPGMA trees in the same manner as sourmash ANI trees. The BUSCO maximum likelihood tree was computed with the same method described above. Processing times and memory usage were tracked using GNU time (v. 1.9) and all code ran on a dual AMD EPYC 7662 64-Core Processor using 32 CPUs.

### Sourmash ANI for fungal taxonomy

To assess the potential for k-mer-based whole genome similarity trees to determine accurate taxonomic placement for fungi and oomycetes, dendrograms were produced from sourmash-estimated ANI matrices and compared to trees produced using more conventional phylogenomic and/or taxonomic inference methods. Taxonomic groups were selected for this analysis to cover a broad range of phyla and include relevant medical, animal and plant pathogens. Groups were selected to consider a number of interesting cases in fungal and oomycete genomics including species complexes, variant/lineage level datasets, high transposable element (TE) content, accessory chromosomes, and wide ranges of genome sizes. Six datasets were constructed spanning genus to order ranks with genome assemblies of fungal and oomycete isolates to consider species rank classification and concordance between sourmash ANI dendrograms and BUSCO maximum likelihood trees.

A range of fungal and oomycete taxa were included to evaluate classification at the species rank. *Aphanomyces* and several relatives in the Saprolegniales were selected as a plant and animal pathogenic oomycete group with genomes available in the NCBI assembly database with some variation in genome sizes and several assemblies with low BUSCO completeness (genome sizes ranging from 39.1 to 104.5 Mbp and single copy BUSCO percentages of 53-57% for three isolates and the rest at 84% or higher). Several other oomycete groups with important plant pathogens, *Phytophthora* clades 1 and 8, were also used. Other selected clades include human pathogenic ascomycetes in the Onygenales *Coccidioides* and *Paracoccidioides*, the basidiomycetous plant pathogenic family known to have high transposable element content Pucciniaceae including *Puccinia* and *Uromyces*, and the rhizosphere-inhabiting Mucoromycota fungi in the Mortierellaceae family including *Mortierella, Linnemannia, Podila, Lunasporangiospora,* and *Gryganskiella*. EarlGrey (v. 4.2.4) was used to assess the extent of transposable element content in Pucciniaceae assemblies.

To consider sourmash ANI tree performance for resolving species complexes, three datasets were constructed considering fungi with known species complexes. Species complexes were chosen from the genera *Rhizopus* and *Mucor* in the order of the Mucorales as representatives from the Mucoromycota phylum of fungi focusing on human and plant pathogens and saprobes, including the *Rhizopus microsporus* and *R*. *arrhizus* complexes. A subset of *Cryptococcus* genomes available in the NCBI assembly database was selected as human pathogenic members of the Basidiomycota, including members of the *C. neoformans* and *C. gattii* complexes, and using *Tremella fuciformis* as an outgroup. *Boletus edulis* was selected as another species complex within the Basidiomycota. A large dataset of 254 genome assemblies from 7 lineages across the northern hemisphere (including assemblies of *Boletus reticulatus* and *B. reticuloceps* as outgroups) was received from Tremble et al. [47], the authors of a study analyzing genomics of this complex, and a summary coalescent gene tree was received from these authors as well for comparison to whole genome similarity trees. The *B. edulis* dataset had a very large range of genome sizes ranging from 20.1 to 349.4 Mbps and a large range of BUSCO completeness percentages ranging from 3.1 to 98.0% [47]. Sourmash ANI trees were compared to BUSCO gene trees and a coalescent orthologous gene tree. *Rhizopus* and *Mucor* genome assemblies in the NCBI assembly database were downloaded with the NCBI dataset function. Select *R. microsporus* genomic reads were downloaded from the NCBI SRA database to have more isolates with variety designations within the *R. microsporus* complex, and genome assemblies were produced as described prior but using the ‘--isolate’ flag in SPAdes. For most BUSCO gene annotations in the species and species complex taxonomic comparisons, the fungi_odb10 database was used for fungi, and stramenopile_odb10 database was used for oomycetes. The only exception was the mucorales_odb12 database was used for the *Rhizopus/Mucor* analysis to ensure a high count of single copy orthologs.

To evaluate the ability of sourmash ANI to resolve subspecies designations, four datasets were used to compare sourmash trees to SNP trees and f3-outgroup dendrograms. As RF distances were only computed at family, genus, and species ranks in the prior analyses, RF distances between subspecies level trees and reference trees were assessed. *Aspergillus fumigatus* was selected as an important human pathogen with intraspecific genetic groups defined by Lofgren et al. [48], *Aspergillus fumigatus* read files available in BioProject PRJNA666940 as well as the NCBI SRA database were assembled. A SNP tree was downloaded from the project’s github repository (https://github.com/MycoPunk/Afum_PopPan) and was pruned to retain isolates that could be assembled and for which subspecific clades_were able to be determined. *Magnaporthe oryzae* was selected as a plant pathogenic subspecies dataset containing some isolates known to contain accessory chromosomes [46] and with an f3-outgroup dendrogram provided by Latorre et al. [49]. For *Magnaporthe oryzae,* 88 assemblies were directly downloaded from BioProject PRJNA354675, and single and paired-end Illumina reads were downloaded for 43 isolates from BioProject PRJNA417903. Reads were assembled as described above. Two isolates were excluded from the *M. oryzae* dataset, SRR6384801 as its reads file contained no quality scores and SRR6384792 due to its low read coverage and BUSCO completeness (43% compared to 98% and higher for most assemblies in the dataset) relative to the rest of the assemblies, and these isolates were pruned from the f3-outgroup dendrogram as well. *Batrachochytrium dendrobatidis* was selected as a member of the relatively undersampled phylum Chytridiomycota and as an important global pandemic pathogen of frogs, and a SNP tree was downloaded from the project’s microreact webpage (https://microreact.org/project/GlobalBd) [50] and was pruned to isolates which could be assembled and for which lineage designations could be determined. Genomic read files for *B. dendrobatidis* were downloaded from NCBI BioProject PRJNA413876.

*Phytophthora ramorum* was selected as an oomycete and important plant pathogen with four well-characterized lineages, and a biallelic SNP-based tree (with runs of homozygosity) was received from the Dale et al. [51]. All sequencing reads files were downloaded from NCBI BioProject PRJNA427329 and assembled using the download and assembly workflow described in the methods. This dataset had very poor BUSCO completeness scores, ranging from 25.9% to 68.4% with a mean of 42.7%. For each subspecies dataset, all sourmash and reference trees, assembly accessions and other identifying information, and, if necessary, isolate renaming csv files are noted in the project’s GitHub and Zenodo repositories: https://github.com/Uehling-Lab/SourmashForFungalTaxonomy/tree/main/SupplementalData_Accessions_Trees_And_BUSCOdatabases/CompareSourmashAndPhylogeneticAnalysis/SubspeciesLevel (DOI: 10.5281/zenodo.21707559). Throughout all comparisons of conventional phylogenomic methods and sourmash ANI trees, all assembly’s NCBI accessions or other identifying information can be found in the project’s GitHub and Zenodo repositories: https://github.com/Uehling-Lab/SourmashForFungalTaxonomy/blob/main/SupplementalData_Accessions_Trees_And_BUSCOdatabases/CompareSourmashAndPhylogeneticAnalysis/Manuscript_Assembly_Metadata.csv (DOI: 10.5281/zenodo.21707559).

## Declarations

### Ethics Approval and Consent to Participate

Not Applicable.

### Consent for Publications

Not Applicable.

### Availability of Data and Materials

The scripts used in analyses as well as relevant data (list of accessions used, tree files, and more) can be found at the SourmashForFungalTaxonomy repository on GitHub and Zenodo (DOI: 10.5281/zenodo.21707559): https://github.com/Uehling-Lab/SourmashForFungalTaxonomy/tree/main

## Acknowledgements

The authors would like to thank Keaton Tremble for providing the *Boletus edulis* assemblies and summary coalescent tree, Angela Dale for providing the SNP-based tree of *Phytophthora ramorum*, and Sergio Latorre for providing the f3-outgroup dendrogram of *Magnaporthe oryzae*.

## Authors’ Contributions

HJ helped design analyses, performed ANI analyses, performed BUSCO tree analysis, interpreted results, and wrote the manuscript. BAV, JKU and NJG obtained funding, designed approaches, interpreted results, and wrote the manuscript. RM and KB contributed to designing approaches and interpreting results. All authors reviewed the manuscript.

