## Supplemental Information for "Whole genome similarity provides a rapid, robust framework for classification of fungal taxa from the genus rank to intraspecies variants"

### Supplement

#### Supplementary Table 1

**Supplementary Table 1** – Summary of publications utilizing whole genome similarity methods for fungal and oomycetous genomic classification.

| DOI (Year) | Broad Whole Genome Similarity Family | Primary Whole Genome Similarity Approach | Primary Taxa Considered | # of Isolates Considered | ANI Usage | Complementary Methods Considered |
|---|---|---|---|---|---|---|
| 10.1016/j.fgb.2025.103969 (2025) | Alignment-based | ANIb (FungANI) | Sordariales (Podospora, Neurospora, Chaetomiaceae) | 123 | Taxonomy + method comparison | Sourmash (Mash distance, k=31) and orthologous gene tree |
| 10.20944/preprints202511.0423.v1 (2025) | Alignment-based | ANIb (FungANI) | Naviculisporaceae (Rhypophila, Pseudorhypophila) | 13 | Taxonomy | BUSCO gene tree |
| 10.1073/pnas.1615148113 (2016) | Alignment-based | ANIb | Rhizopus (Mucoromycota) | 2 | Genome comparison | |
| 10.1371/journal.pone.0144769 (2015) | Alignment-based | ANIb | Rhizoctonia (Cantharellales) | 4 | Genome comparison | |

| | | | | | | |
|---|---|---|---|---|---|---|
| 10.3390/jof8030269 (2022) | Alignment-based | ANIb | Ustilaginaceae | 20 | Taxonomy / species delimitation | Core genome tree and AAI (average amino acid identity) |
| 10.1371/journal.ppat.1011510 (2023) | Alignment-based | OrthoANI | Microsporidia | 15 | Taxonomy | Orthologous gene tree |
| 10.1111/jeu.12944 (2022) | Alignment-based | OrthoANI | Microsporidia | 65 | Taxonomy / species delimitation | Orthologous gene tree and SSU rRNA tree |
| 10.1128/mbio.00582-24 (2024) | Alignment-based | OrthoANI | Microsporidia | 2 | Genome comparison | |
| 10.1007/s10482-020-01480-9 (2020) | Alignment-based | OrthoANI | Yeasts (Metschnikowia) | 71 | Taxonomy / species delimitation | Orthologous gene tree |
| 10.1186/s12864-025-12223-3 (2025) | Alignment-based | OrthoANI | Onygenaceae | 9 | Taxonomy | |
| 10.1371/journal.pone.0210792 (2019) | Alignment-based | OrthoANIu | Yeasts (Hanseniaspora) | 11 | Taxonomy | Multi-locus gene tree |
| 10.1093/femsyr/foaa042 (2020) | Alignment-based | OrthoANIu | Yeasts (broad, Asco and Basidio) | 30 | Taxonomy | dDDH (digital DNA-DNA hybridization) and $K_r$ |
| 10.1371/journal.pgen.1011945 (2025) | Alignment-based | OrthoANIu | Cryptococcus | 39 | Taxonomy / species delimitation | Orthologous gene trees (supermatrix and coalescent) and AAI |
| 10.1073/pnas.2319389122 (2024) | Alignment-based | BUSCO-based ANI | Fungi (broad) | 233 | Species boundary delimitation | |

| | | | | | | |
|---|---|---|---|---|---|---|
| 10.1111/mpp.12765 (2018) | Alignment-based | ANIm | Phytophthora, Notophytophthora (oomycetes) | 12 | Taxonomy / species delimitation | SNP-based tree (REALPHY) |
| 10.1016/j.simyco.2015.07.001 (2015) | Alignment | ANIm | Alternaria | 11 | Taxonomy / species delimitation | SNP-based tree (REALPHY) |
| 10.3390/jof10090646 (2024) | Mapping-based | FastANI + MOAG [a BUSCO-based approach] | Yeasts (broad, Asco and Basidio) | 644 | Species boundary delimitation | |
| 10.3389/fmicb.2023.1199144 (2023) | Mapping-based | FastANI | Monascus (Aspergillaceae) | 15 | Species boundary delimitation | Orthologous gene tree |
| 10.3390/jof11050401 (2025) | Mapping-based | FastANI | Trichosporonales | 16 | Taxonomy | Orthologous gene tree |
| 10.1186/s12864-020-6653-6 (2020) | Mapping-based | FastANI | Trichoderma (Hypocreales) | 10 | Taxonomy / species delimitation | Orthologous gene tree |
| 10.1016/j.cell.2024.04.043 (2024) | Mapping-based | FastANI | Fungi (broad) | 11,623 | Species boundary delimitation | |
| 10.1186/s12864-023-09718-2 (2023) | Mapping-based | FastANI | Cutaneotrichosporon | 5 | Genome comparison | dDDH and ITS percent identity |
| 10.1186/s43008-019-0011-9 (2019) | Mapping-based | FastANI | Tilletia | 10 | Species boundary delimitation | Coalescent gene tree |

| | | | | | | |
|---|---|---|---|---|---|---|
| 10.1186/s12879-025-10696-x (2025) | Not disclosed (alignment or mapping-based) | Undisclosed method using PyANI | Candida | 3 | Genome comparison | |
| 10.3390/jof6040246 (2020) | Sketch | Mash, Dashing (k=12–22) | Fungi (broad) | 1,544 | Species boundary exploration | K-mer overlap |
| 10.1093/nar/gkae609 (2024) | Sketch | GSearch (k=21) | Fungi (broad) | 9,700 | Genome search / similarity | Many sketch-based methods |
| 10.3390/jof9020183 (2023) | K-mer analysis (not disclosed if sketch based) | K-mer tree | Aspergillus | 13 | Taxonomy | Single gene percent identity |

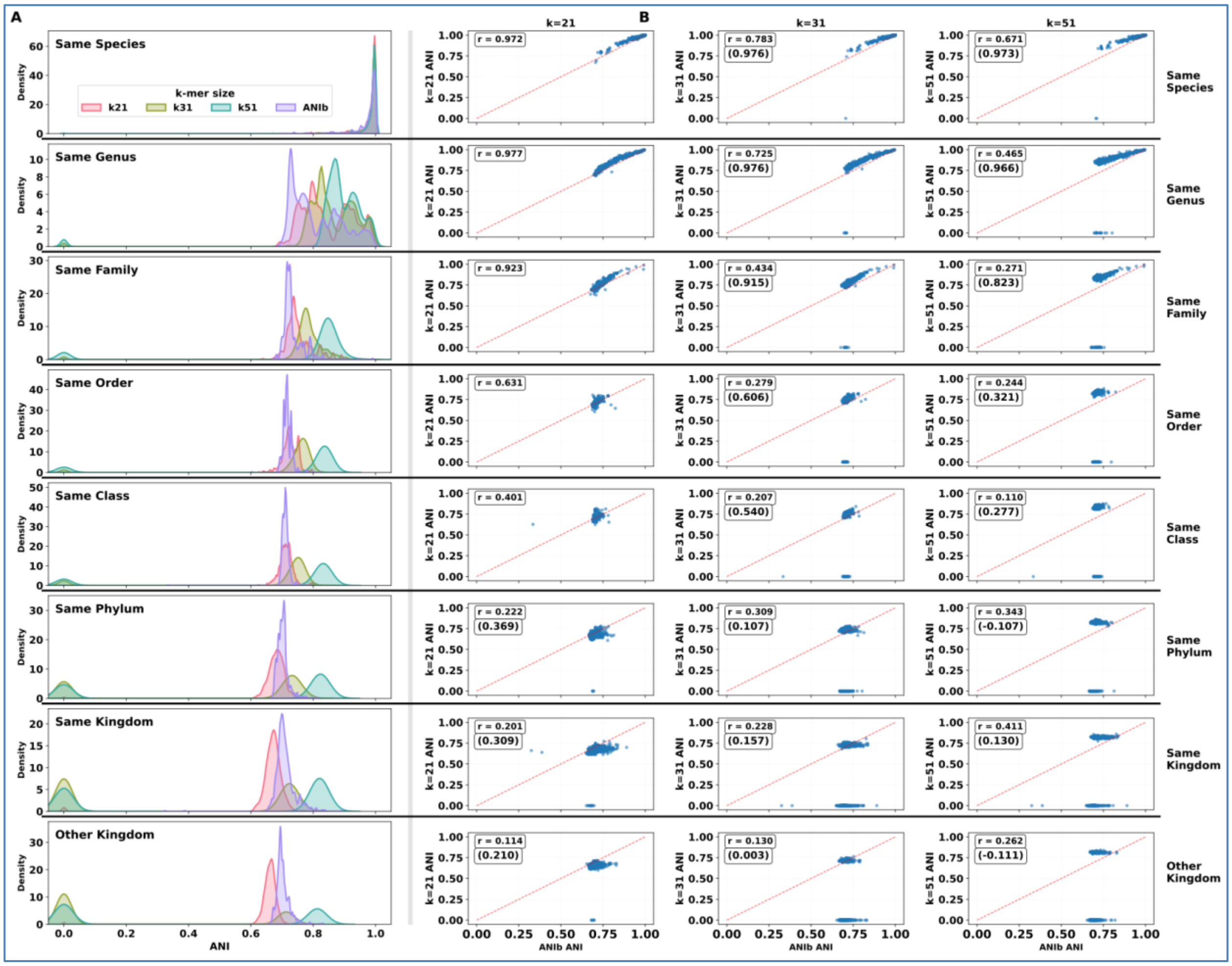

**Supplementary Figure 1** – Comparative similarity analysis of ANI methods computed by ANIb and sourmash Jaccard similarity ANI methods at various k-mer lengths across evolutionary paths. (A) Kernel density estimation plots for ANI as determined with ANIb and Sourmash ANI with k-mer sizes of 21, 31, and 51 using Jaccard similarity. Each row displays the distribution of ANI between all target assemblies and all assemblies in the evolutionary path at the specified taxonomic rank, from same species to same and other kingdoms. (B) Linear regressions between ANIb and sourmash ANI by k-mer size for the same dataset. Rows correspond to the evolutionary paths outlined on the left. Pearson's r values for the full data set are given at the top right, with r values computed excluding sourmash ANI values of 0 in parentheses. In each plot, red dashed lines indicate a slope of 1.0.

**Supplementary Table 2**– ANI distribution statistics for evolutionary path analysis comparing target assemblies to genome assemblies of increasing taxonomic distance using ANIb and sourmash-based Jaccard similarity methods at various k-mer lengths including k21, k31, and k51. Mean and standard deviation of ANI estimates for each rank in the evolutionary paths are displayed, corresponding to the Supplementary Figure 1 KDE plots with 0% values excluded from statistics.

| Category | ANIb | k21 | k31 | k51 |
|---|---|---|---|---|
| **Same Species** | 97.4% ± 4.3% | 98.3% ± 3.3% | 98.5% ± 2.7% | 98.8% ± 2.2% |
| **Same Genus** | 81.1% ± 8.0% | 84.7% ± 7.8% | 86.9% ± 6.5% | 90.0% ± 4.6% |
| **Same Family** | 73.4% ± 3.4% | 75.6% ± 4.6% | 79.1% ± 3.8% | 85.2% ± 2.3% |
| **Same Order** | 71.7% ± 1.4% | 72.6% ± 2.4% | 76.3% ± 2.1% | 83.6% ± 1.2% |
| **Same Class** | 71.1% ± 1.6% | 71.3% ± 2.3% | 75.0% ± 1.8% | 83.3% ± 1.2% |
| **Same Phylum** | 70.2% ± 1.8% | 68.3% ± 2.5% | 73.3% ± 1.6% | 82.4% ± 1.0% |
| **Same Kingdom** | 71.0% ± 3.3% | 67.2% ± 2.3% | 72.5% ± 1.4% | 82.0% ± 0.9% |
| **Other Kingdom** | 70.3% ± 2.1% | 66.2% ± 1.7% | 71.4% ± 1.1% | 81.3% ± 0.6% |

**Supplementary Table 3** – Correlation between between ANIb similarity values and sourmash ANI values determined using either Jaccard similarity or max containment using k-mer sizes k21, k31, and k51 for the evolutionary path analysis comparing target assemblies to genome assemblies of increasing taxonomic distance. Linear regression correlation coefficients between ANIb and sourmash ANI at each k-mer sie are displayed, and the correlation coefficients computed without considered ANI values of 0% are shown in parenthesis.

| | **Jaccard k21** | **Jaccard k31** | **Jaccard k51** | **Max Containment k21** | **Max Containment k31** | **Max Containment k51** |
|---|---|---|---|---|---|---|
| **Same Species** | 0.97 | 0.78 (0.98) | 0.67 (0.97) | 0.97 | 0.77 (0.97) | 0.67 (0.97) |
| **Same Genus** | 0.98 | 0.73 (0.98) | 0.47 (0.97) | 0.98 | 0.73 (0.98) | 0.47 (0.97) |
| **Same Family** | 0.92 | 0.43 (0.92) | 0.27 (0.82) | 0.93 | 0.43 (0.92) | 0.27 (0.83) |
| **Same Order** | 0.63 | 0.28 (0.61) | 0.24 (0.32) | 0.64 | 0.28 (0.62) | 0.24 (0.34) |
| **Same Class** | 0.40 | 0.21 (0.54) | 0.11 (0.28) | 0.32 | 0.21 (0.52) | 0.11 (0.27) |
| **Same Phylum** | 0.22 (0.34) | 0.31 (0.11) | 0.35 (-0.11) | 0.17 (0.30) | 0.31 (0.07) | 0.34 (-0.17) |
| **Same Kingdom** | 0.20 (0.31) | 0.23 (0.16) | 0.42 (0.13) | 0.18 (0.23) | 0.23 (0.16) | 0.41 (0.13) |
| **Different Kingdom** | 0.11 (0.21) | 0.13 (0.00) | 0.26 (-0.11) | 0.09 (0.13) | 0.13 0.01) | 0.26 (-0.09) |

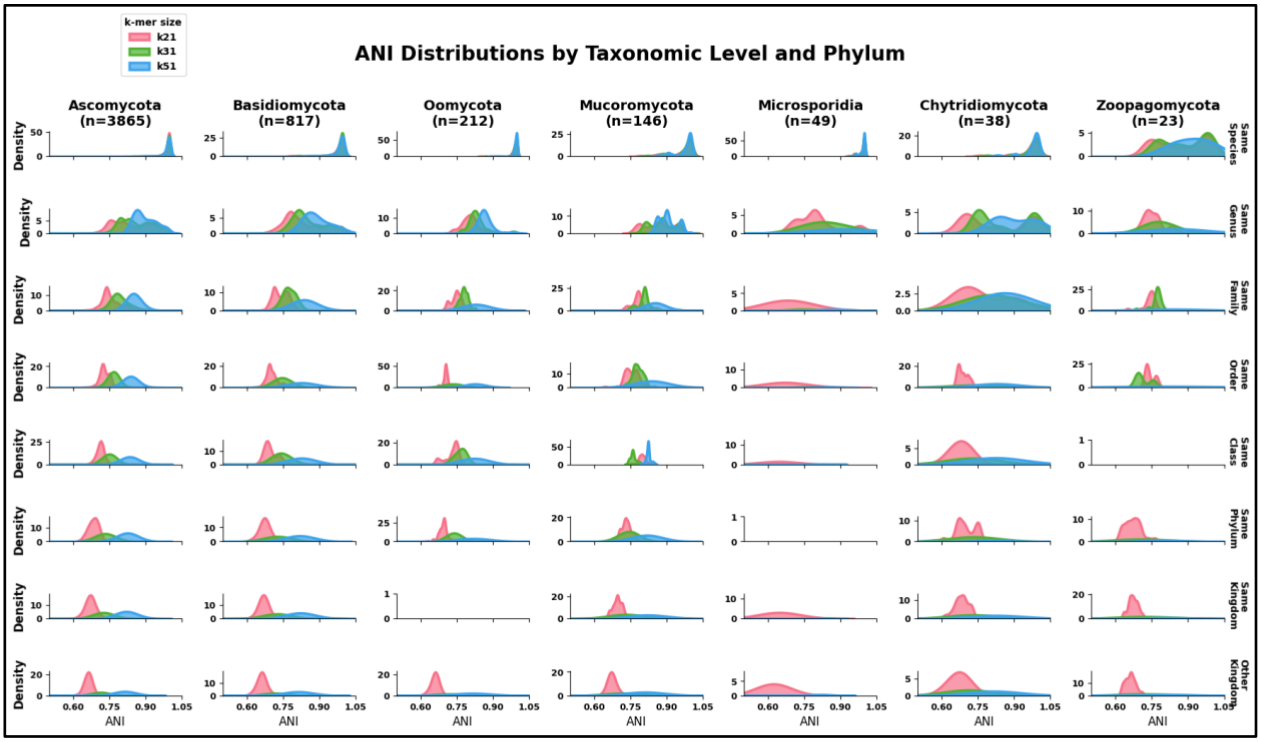

**Supplementary Figure 2** – Kernel density estimate plots of sourmash Jaccard similarity derived ANI values at 3 k-mer sizes (k21, k31, and k51) broken down by the top 7 phyla by number of evolutionary paths. The x-axis for each plot is scaled from 0.55 to 1.05 ANI which leaves out the ANI values of zero, but makes it easier to visually assess the higher ANI range.

**Supplementary Table 4** – BUSCO database and number of single copy BUSCO genes used to produce the phylogenetic trees used as references in the sourmash parameter determination analysis comparing k-mer sizes (16, 21, 31, 51, 61, and 71) and ANI estimation methods (Jaccard similarity and max containment). The phylogenetic trees are found in Figure 2.

| Family | SameSpecies | SameGenus | SameFamily | BUSCO Database |
|---|---|---|---|---|
| **Cordycipitaceae** | 677 | 644 | 259 | fungi_odb10 |
| **Boletaceae** | 268 | 222 | 345 | fungi_odb10 |
| **Hypocreaceae** | 621 | 713 | 720 | fungi_odb10 |
| **Lichtheimiaceae** | 390 | 550 | 278 | fungi_odb10 |
| **Mortierellaceae** | 566 | 550 | 363 | fungi_odb10 |
| **Mucoraceae** | 209 | 283 | 476 | fungi_odb10 |
| **Nectriaceae** | 751 | 707 | 690 | fungi_odb10 |
| **Peronosporaceae** | 841 | 866 | 1427 | oomycota_odb12 |
| **Pleosporaceae** | 674 | 687 | 676 | fungi_odb10 |
| **Polyporaceae** | 603 | 456 | 555 | fungi_odb10 |
| **Pythiaceae** | 1942 | 1205 | 1653 | oomycota_odb12 |
| **Trichosporonaceae** | 649 | 659 | 650 | fungi_odb10 |
| **Ustilaginaceae** | 731 | - | 690 | fungi_odb10 |

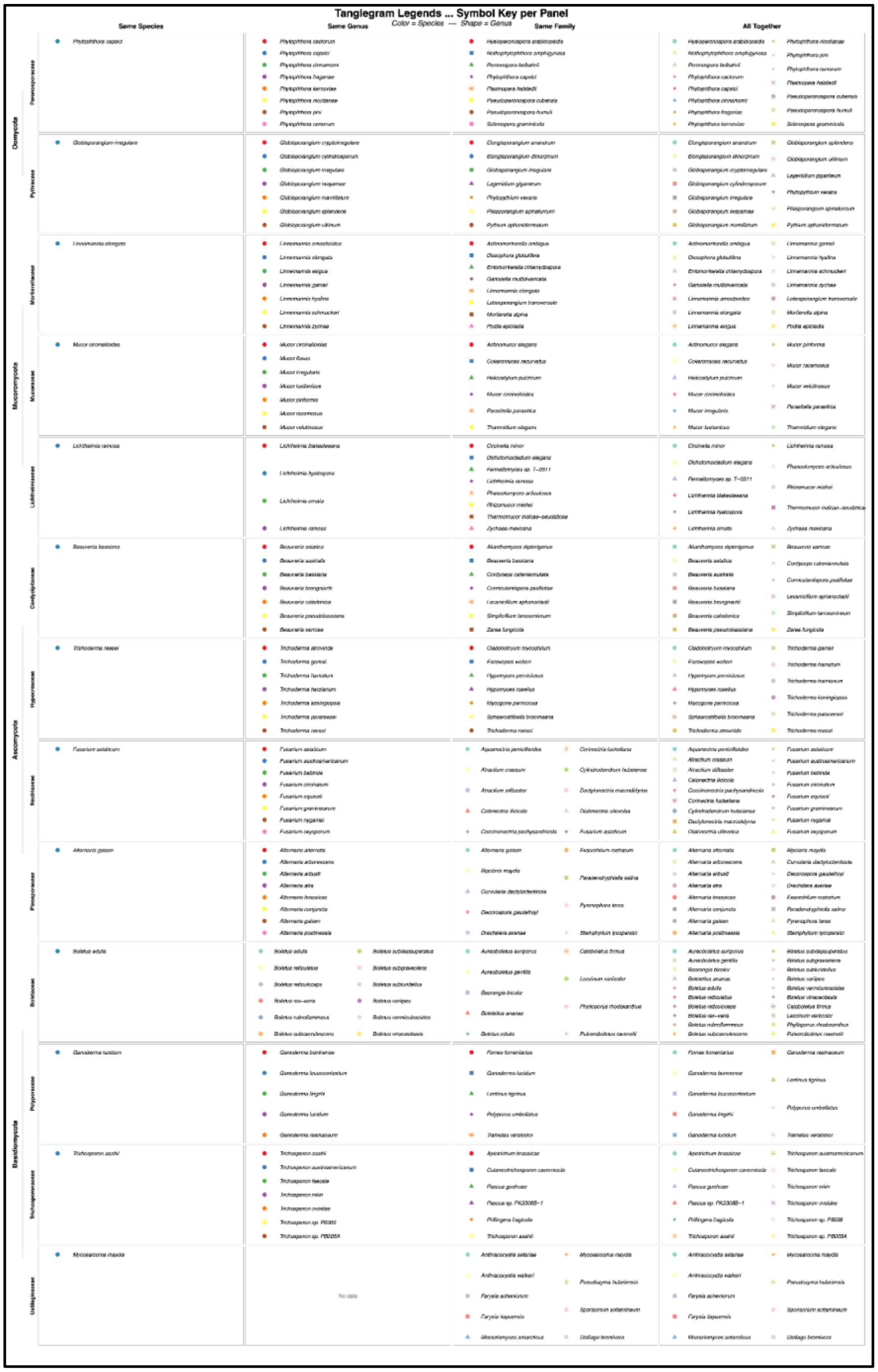

**Supplementary Figure 3** – Legend denoting the species each symbol refers to in the Figure 2 tanglegrams displaying concordance between sourmash ANI-based trees and concatenated BUSCO gene maximum likelihood phylogenetic trees.

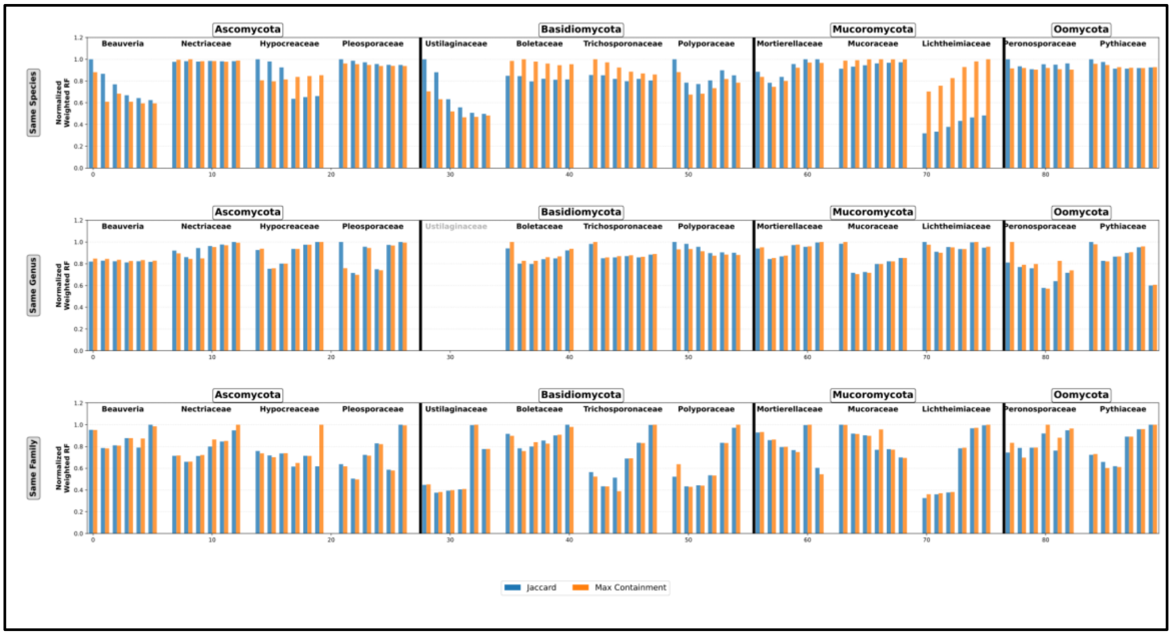

**Supplementary Figure 4** – Weighted Robinson-Foulds (RF) distances denoting similarity between sourmash ANI-trees and concatenated single copy BUSCO gene maximum likelihood trees for a broad range of fungi and oomycete taxa at the same species, same genus, and same family ranks. RF distances were computed between BUSCO trees and sourmash-ANI trees created with a variety of k-mer sizes (16, 21, 31, 51, 61, and 71) and two similarity estimation methods (Jaccard similarity and max containment), were normalized to a range between 0 and 1, and the sourmash methods with the lowest RF distances were considered to create trees most similar to the reference BUSCO trees.

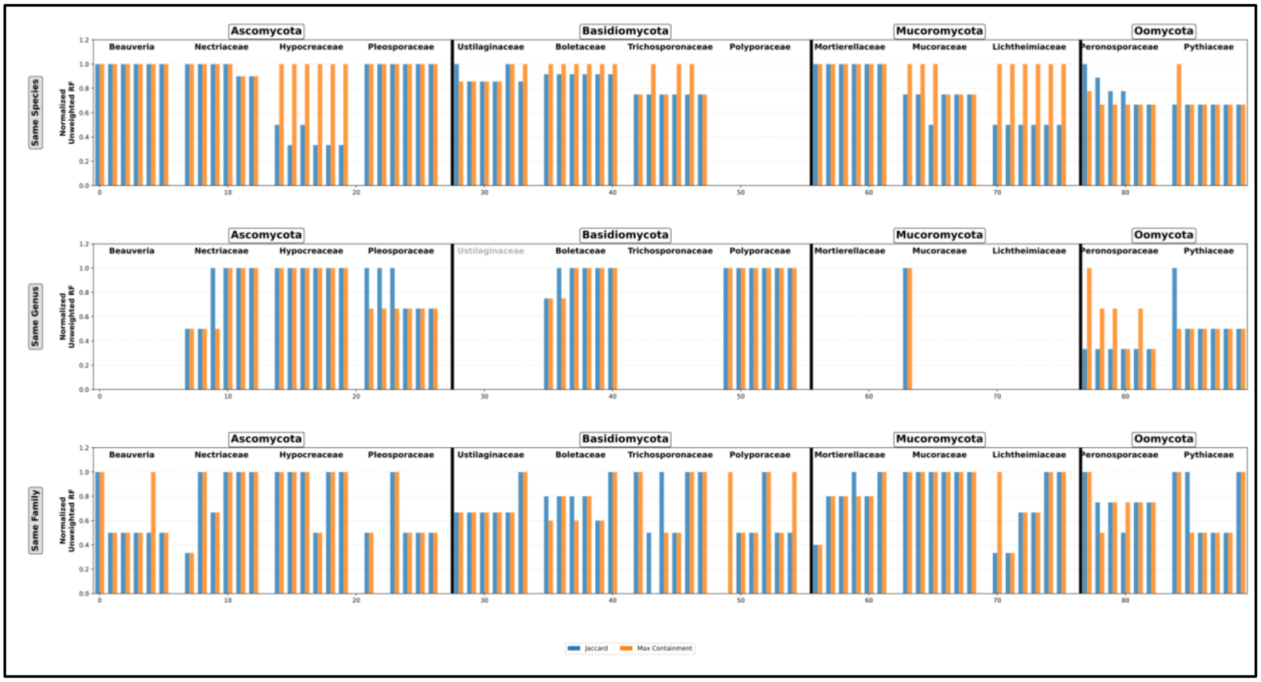

**Supplementary Figure 5** – Unweighted Robinson-Foulds (RF) distances denoting similarity between sourmash ANI-trees and concatenated single copy BUSCO gene maximum likelihood trees for a broad range of fungi and oomycete taxa at the same species, same genus, and same family ranks. RF distances were computed between BUSCO trees and sourmash-ANI trees created with a variety of k-mer sizes (16, 21, 31, 51, 61, and 71) and two similarity estimation methods (Jaccard similarity and max containment), were normalized to a range between 0 and 1, and the sourmash methods with the lowest RF distances were considered to create trees most similar to the reference BUSCO trees.

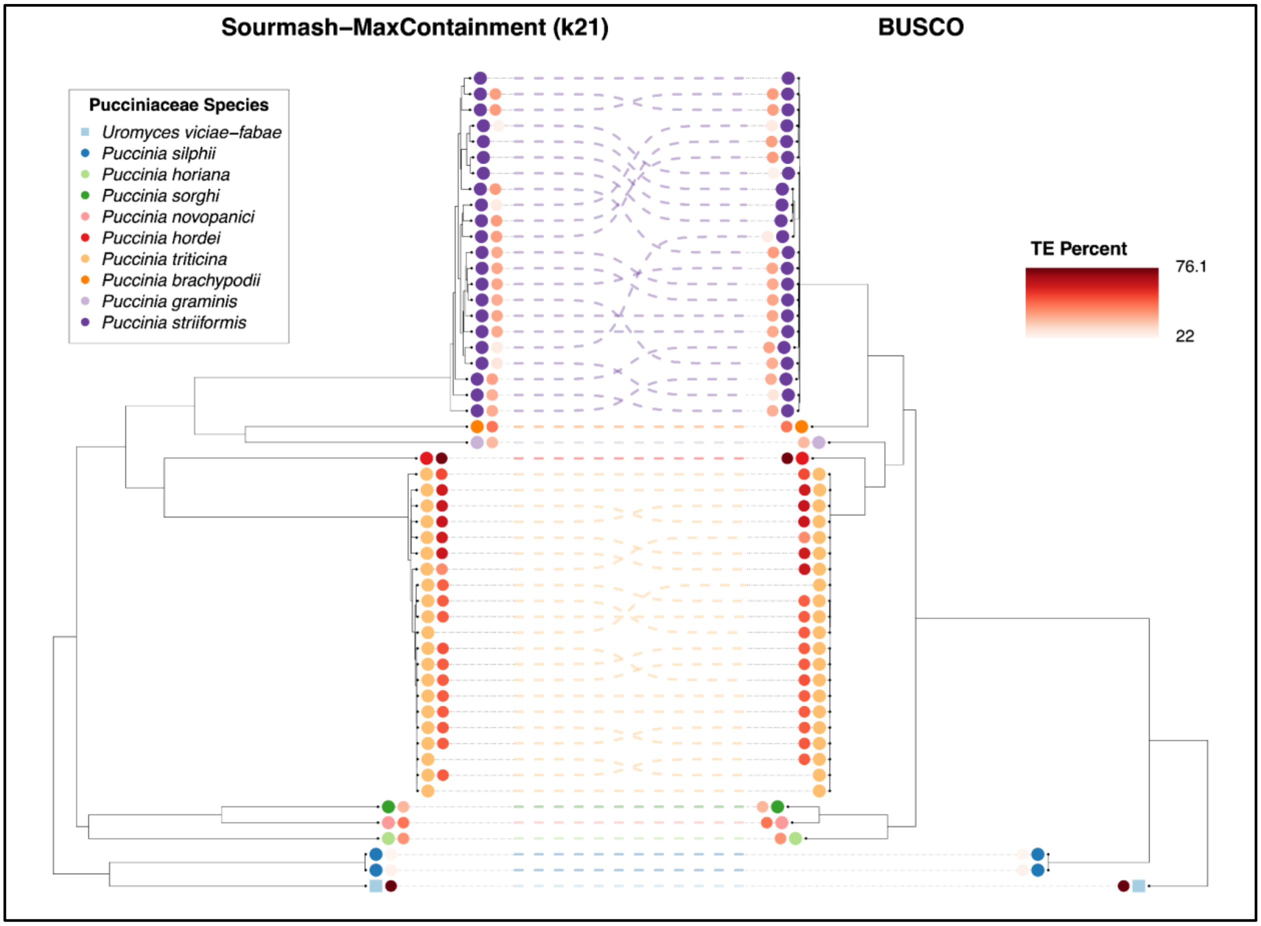

**Supplementary Figure 6** – Concordance between a sourmash k21 max containment ANI tree (left side tree) and a BUSCO gene maximum likelihood tree (right side tree) displaying clade agreement between species in the Pucciniaceae with total transposable element percentage noted as estimated using EarlGrey.

| k-mer Size | Dataset | Misclassification Count | |
|---|---|---|---|
| | | Jaccard | Max Containment |
| k16 | Boletus edulis | 11 | 1 |
| k21 | Boletus edulis | 9 | 3 |
| k31 | Boletus edulis | 4 | 2 |
| k51 | Boletus edulis | 4 | 1 |
| k61 | Boletus edulis | 3 | 1 |
| k71 | Boletus edulis | 3 | 1 |
| k16 | Aspergillus fumigatus | 14 | 4 |
| k21 | Aspergillus fumigatus | 13 | 18 |
| k31 | Aspergillus fumigatus | 13 | 1 |
| k51 | Aspergillus fumigatus | 9 | 0 |
| k61 | Aspergillus fumigatus | 6 | 0 |
| k71 | Aspergillus fumigatus | 5 | 0 |
| k16 | Magnaporthe oryzae | 5 | 6 |
| k21 | Magnaporthe oryzae | 13 | 6 |
| k31 | Magnaporthe oryzae | 10 | 12 |
| k51 | Magnaporthe oryzae | 13 | 13 |
| k61 | Magnaporthe oryzae | 5 | 12 |
| k71 | Magnaporthe oryzae | 5 | 12 |
| k16 | Batrachochytrium dendrobatidis | 1 | 0 |
| k21 | Batrachochytrium dendrobatidis | 1 | 0 |
| k31 | Batrachochytrium dendrobatidis | 1 | 0 |
| k51 | Batrachochytrium dendrobatidis | 0 | 0 |
| k61 | Batrachochytrium dendrobatidis | 0 | 0 |
| k71 | Batrachochytrium dendrobatidis | 0 | 0 |
| k16 | Phytophthora ramorum | 1 | 0 |
| k21 | Phytophthora ramorum | 1 | 0 |
| k31 | Phytophthora ramorum | 1 | 0 |
| k51 | Phytophthora ramorum | 1 | 0 |
| k61 | Phytophthora ramorum | 1 | 0 |
| k71 | Phytophthora ramorum | 1 | 0 |

**Supplementary Table 5 –** Number of mis-classifications per sourmash k-mer size and similarity estimation method combination for taxa with species complex or subspecific lineage level datasets.

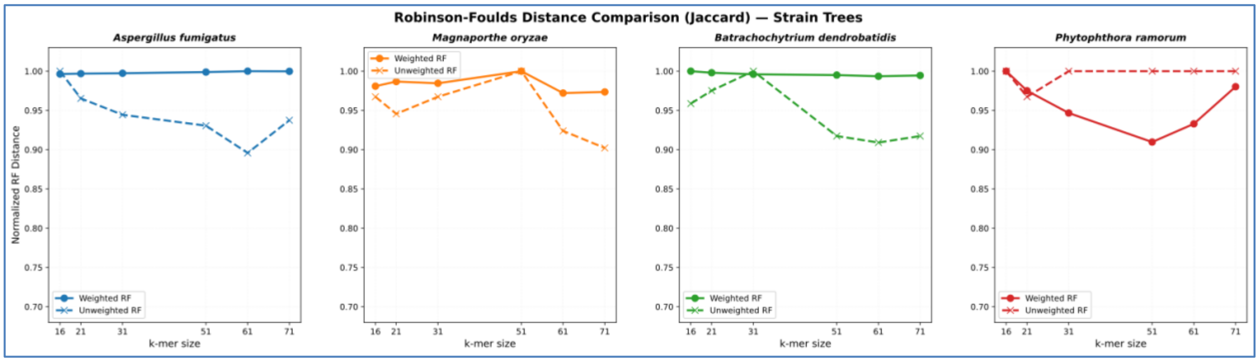

**Supplementary Figure 7** – Normalized RF distances computed between SNP-based trees and Jaccard-based sourmash ANI trees at various k-mer sizes from k16 to k71.

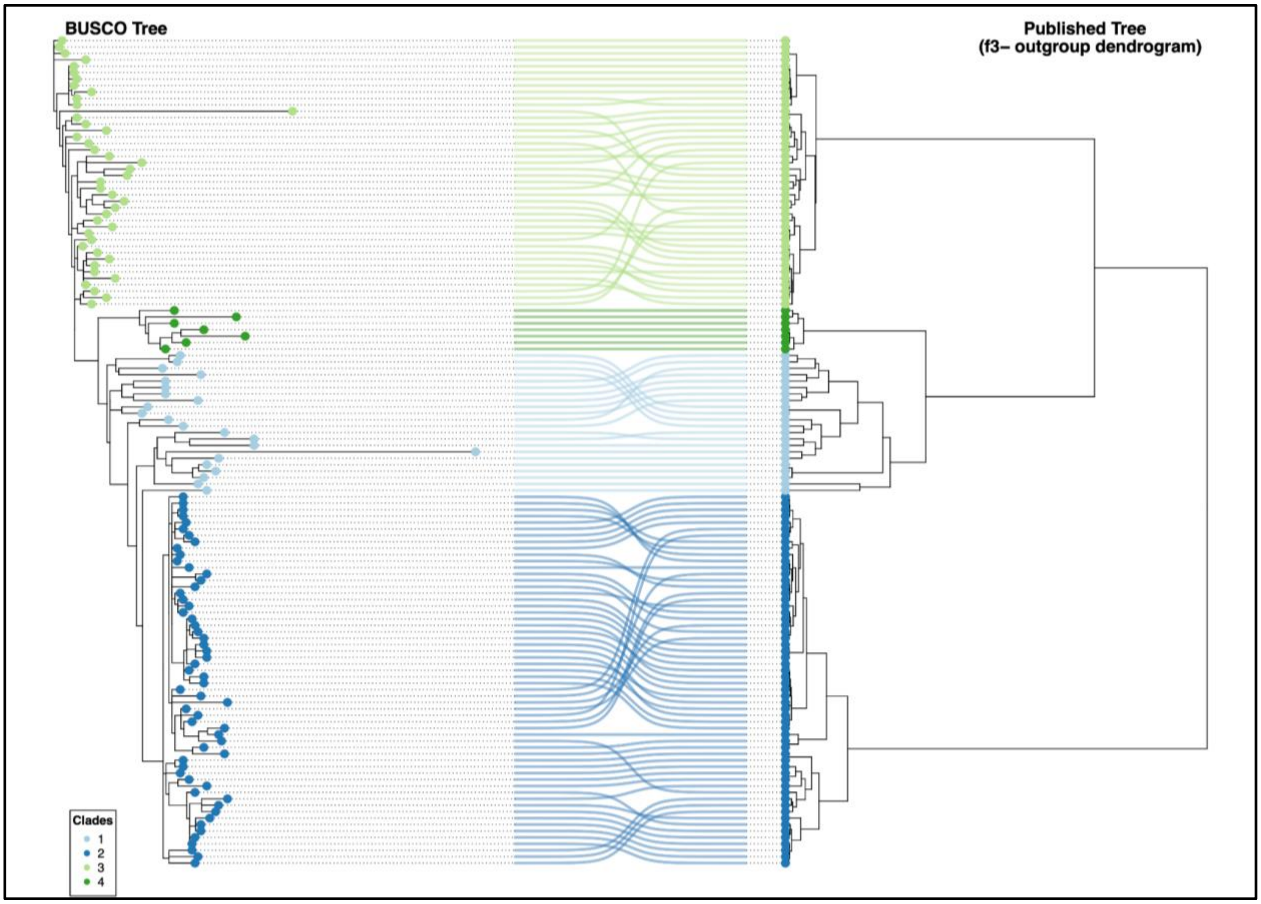

**Supplementary Figure 8** – Concordance between a 580 BUSCO gene maximum likelihood tree (left side tree) and an f3-outgroup dendrogram (right side tree) displaying clade agreement between subspecific clades of *Magnaporthe oryzae* .

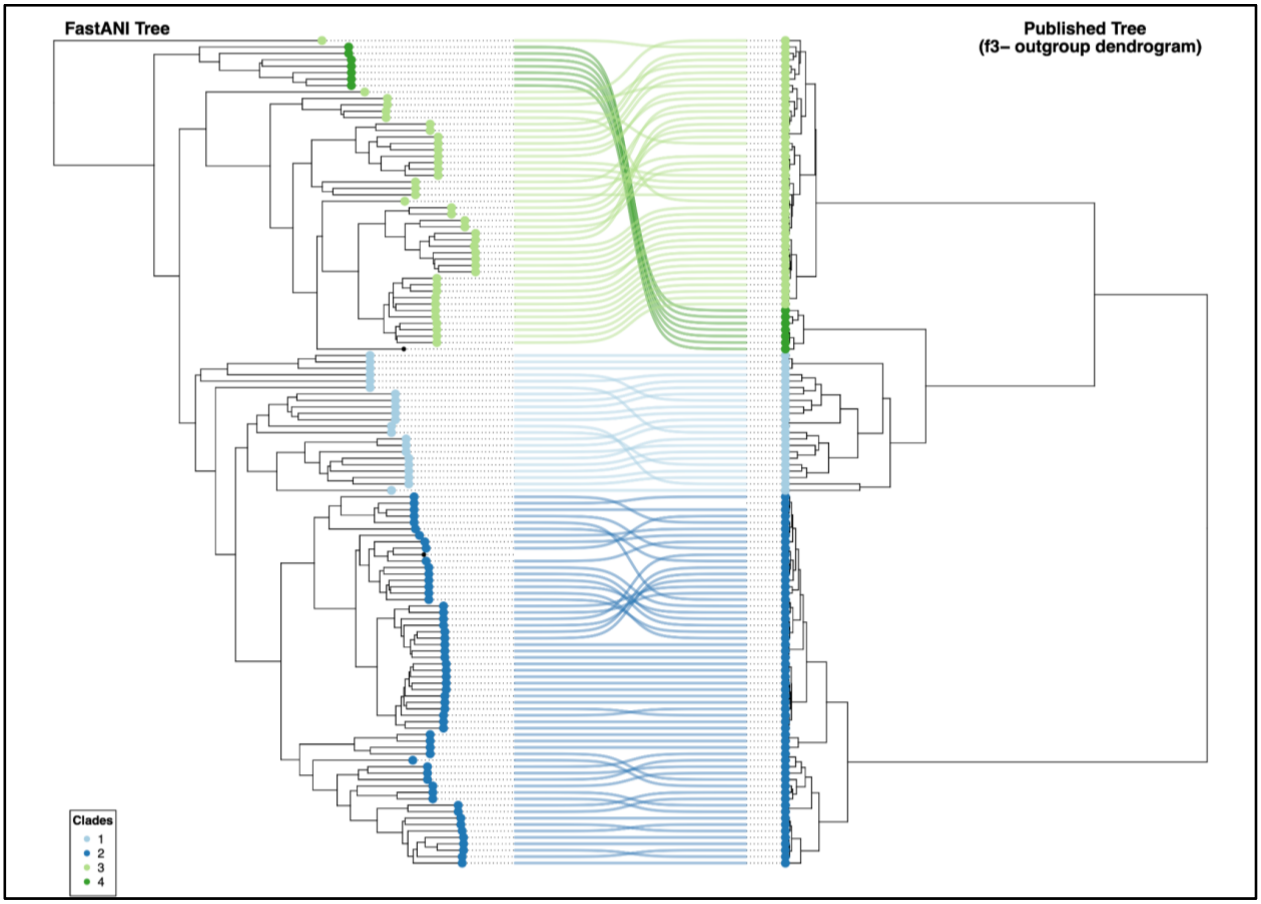

**Supplementary Figure 9** – Concordance between a FastANI-based whole genome similarity tree (left side tree) and an f3-outgroup dendrogram (right side tree) displaying clade agreement between subspecific clades of *Magnaporthe oryzae*. FastANI was computed as reciprocal ANI using pyANI.

### Supplemental Text

#### Supplemental Results

To compare the performance of sourmash-inferred ANI and ANIb methods for estimating genome similarity of fungi and oomycetes across a broad range of taxonomic distances, assemblies were compared to an array of genomes of increasing evolutionary distance. ANI was quantified between target genome assemblies and those sharing the same genus (interspecific), family (intergeneric), order (interfamilial), class (interorder), phylum (interclass), and kingdom (interphylum), as well as assemblies from a different kingdom (with oomycetes compared to fungi and vice versa). Similar to Pierce-Ward et al. [52], we refer to these sets of genome assemblies at increasing evolutionary distances as 'evolutionary paths' throughout this manuscript. There were 5,261 evolutionary paths created from 16,228 unique genome assemblies in the NCBI Assembly database with 1,754 target species belonging to the following phyla: Ascomycota (n=3,865), Basidiomycota (n=817), Oomycota (n=212), Mucoromycota (n=146), Microsporidia (n=49), Chytridiomycota (n=38), unknown (n=22), Blastocladiomycota (n=3), and Cryptomycota (n=1). A csv file with all evolutionary paths can be found at the project's GitHub and Zenodo repositories: https://github.com/Uehling-Lab/SourmashForFungalTaxonomy/tree/main/SupplementalData_Accessions_Trees_And_BUSCOdatabases/EvolutionaryPathAnalysis (DOI: 10.5281/zenodo.21707559). Sourmash was used for all 5,261 evolutionary paths corresponding to 38,171 pairwise comparisons. Due to its considerably slower processing, ANIb was only used for 1,056 evolutionary paths.

We compared ANIb to sourmash's ANI at three different k-mer sizes (k = 21, 31, and 51). ANI values of 0% were not considered in these statistics given they result from sourmash not having sufficient k-mer matches to produce an ANI estimation. Same species comparisons for ANIb and all k-mer sizes of sourmash ANI produced distributions of whole genome similarity in a tight range with means around 98% and standard deviations around 2-4%, while comparing genomes of target species to intrageneric, heterospecific genome assemblies resulted in a wide ANI range for all methods with means and standard deviations of 81 ± 8%, 85 ± 8%, 86 ± 10%, and 88 ± 14% for ANIb, k21, k31, and k51, respectively (Supplementary Figure 1A, Supplementary Table 2). As evolutionary paths increased in taxonomic distance from intrafamily up to kingdom ranks comparisons, all three sourmash, Jaccard-derived ANI distributions saw a slight, consistent decrease in mean ANI values, and the degree of decrease in mean ANI was inversely correlated with k-mer size (Supplementary Figure 1A). Sourmash, especially larger k-mer sizes of 31 and 51, estimated many ANI values to be 0% for larger taxonomic distances.

Correlations between sourmash ANI and ANIb values from the evolutionary path analyses were compared using linear regression (Supplementary Table 3). Linear regression plots between Jaccard-inferred ANI and ANIb are shown in Supplementary Figure 1B. ANIb correlates highly with k21 for same species, same genus, and same family rank comparisons with correlation coefficients of 0.97, 0.98, and 0.92 respectively. At greater taxonomic distances the agreement declines rapidly between

ANIb and sourmash ANI as most ANI methods are not trustworthy below 75% [36] and are not recommended for use below the 75-80% ANI range [35] (Supplementary Figure 1B). At the same species through the same family ranks, both k31 and k51 show less agreement with ANIb overall when ANI values of 0% are considered, but when these values are excluded from the analysis the regression coefficients more closely match the patterns seen for k21. Using a k-mer size of 51 led to a quicker breakdown of correlation with ANIb as taxonomic distance increased. These results highlight agreement between ANIb and sourmash ANI estimation, especially using a k-mer size of 21. Sourmash was thus used for the remaining analysis given its correlation with ANIb and due to its speed.

Next, distributions of whole genome similarity for k21, k31, and k51 were assessed on a per phylum basis. A KDE figure for ANI values, determined using Jaccard similarity, can be found in Supplementary Figure 1. The top 7 phyla by number of evolutionary paths are shown, leaving out evolutionary paths for the isolates of unknown phyla due to unresolved placement, as well as Blastocladiomycota and Cryptomycota due to their low count of evolutionary paths.

The Ascomycota, Basidiomycota, and Oomycota displayed much the same trends in ANI distributions as the overall dataset in Supplementary Figure 1 with a tight intraspecific ANI distribution of means and standard deviations of 98 ± 4%, 97 ± 5%, and 99 ± 2% at k21 respectively and decreasing mean ANI with increasing taxonomic distance (Supplementary Figure 1). Microsporidia have a similarly tight intraspecies ANI range at k21 of 99 ± 2% but broader ANI ranges for intrafamilial and higher rank comparisons (k21 intrafamilial standard deviation of 28% compared to 8%, 8%, and 6%

for Ascomycota, Basidiomycota, and Oomycota respectively). Mucoromycota and Chytridiomycota follow the overall Supplementary Figure 1 trends to a lesser degree, displaying a relatively tight range of same species ANI values with tails extending past 90% ANI, and with Chytridiomycota displaying a broader range of intrageneric and intrafamilial k21 ANI values (13% and 20% respectively) compared to the more strongly represented taxa mentioned prior (Supplementary Figure 1). For the Zoopagomycota, same species ANI values encompassed a much broader range than all other phyla at the intraspecific rank with a mean and standard deviation of 87 ± 10% at k21. While the ANI distributions observed in Ascomycota, Basidiomycota, and Oomycota are highly similar, other fungal phyla display varied ANI distributions.

#### Supplemental Discussion

Comparing ANI computed by sourmash methods to ANIb revealed that the methods broadly produce similar ANI distributions for pairwise genome assembly comparisons between isolates in the same family and at lower taxonomic ranks, especially for k21 but also for larger k-mer sizes where non-zero ANI can be estimated. This work noted a mean ANI within species near 98% for all ANI methods assessed with a standard deviation near 3%; although, as seen in other analysis in this work, leveraging NCBI taxonomic assignments comes with the risk of misidentifications which would lead to inaccuracies in our within species ANI estimates. The intraspecific ANI distribution we report using sourmash is similar to what other authors reported using non-sketch-based ANI methods. A study applying FastANI to yeast genomes reports within-species ANI cutoffs ranging from 92 to 98% [24] and a broader study reports a species cutoff of 95%. Studies using BLAST-based methods like ANIb and OrthoANI report similar findings like most Microsporidia isolates in one study having intraspecific ANI above 95% with some outliers, and a study in the Sordariales recommends a species cutoff at 99% [9] . Moran et al. used a core genome ANI method and found average minimum intraspecific ANI of 98.5% and 97.6% for Ascomycetes and Basidiomycetes respectively but also notes a very broad range of minimum intraspecific ANI from 85.8% to over 99.9% [22]. The wide range of ANI values from different studies likely reflects the different ANI methods used, the variety of taxa considered, the number of assemblies considered, and potential mis-identified isolates and cryptic species which can increase or decrease estimates. An ANI of 95% is considered, with exceptions, to be a generally reliable speciation threshold in bacteria [26], and, given the range of ANI values reported for fungal intraspecific ANI, future studies are required to confirm if a speciation threshold or thresholds exist for fungi and oomycetes. We hypothesize that the discrepancy in ANI KDE distributions between more densely sampled phyla like Ascomycota and Basidiomycota as compared to Chytridiomycota and Zoopagomycota likely reflects the more mature phylogenomic characterization and broader sampling of the Dikarya rather than strictly biological reality in these relatively understudied groups.

#### Supplemental Methods

All fungal and oomycetous assemblies in NCBI were downloaded using the ‘--exclude-atypical’ flag, which excludes assemblies flagged for a variety of reasons including chimeras, contamination, and invalid genome sizes [53], and ANI was quantified between target genome assemblies and assemblies sharing the same genus (interspecific), family (intergeneric), order (interfamilial), class (interorder), phylum (interclass), and kingdom (interphylum), as well as assemblies from a different kingdom (with oomycetes compared to fungi and vice versa). Each evolutionary path is anchored by a target assembly, with target assemblies chosen by considering all species with more than one genome assembly to facilitate intraspecific comparisons, but no more than 10 isolates per species were considered target assemblies to limit overrepresentation. To construct evolutionary paths, 11 genome assemblies from the same species as the target assembly, with 10 randomly designated as target assemblies, were paired sequentially (e.g., assembly 1 compared to 2, 2 to 3, and so on), whereas assemblies representing higher taxonomic ranks were randomly selected from all genome assemblies to satisfy the defined rank relationships (same genus, same family, etc.). A list of assemblies used in this analysis can be found in the project's GitHub and Zenodo repositories: https://github.com/Uehling-Lab/SourmashForFungalTaxonomy/blob/main/SupplementalData_Accessions_Trees_And_BUSCOdatabases/FamilyGenusSpecies_SourmashANIParameterAnalysis/Assembly_Taxonomy_Matrix.csv (DOI: 10.5281/zenodo.21707559).

Sourmash was used to determine ANI for all pairwise comparisons defined by the evolutionary paths. For each evolutionary path, the sourmash sketch function was used to compute sketches for all genome assemblies using k-mer sizes of 21, 31, and 51 bp (referred to as 'k21', 'k31', and 'k51' throughout) followed by comprehensive pairwise ANI computation using the sourmash compare function. ANIb was also assessed on the evolutionary paths but was only computed on a subset of the evolutionary paths due to much longer processing times. ANIb was computed using the 'average_nucleotide_identity.py' script of pyani (v 0.2.12), and ANI reported for ANIb is the mean ANI of reciprocal pairwise comparisons. To assess results, kernel density estimation (KDE) plots (Jaccard similarity only) and linear regression plots were produced in python using the libraries seaborn (v. 0.13.2) and matplotlib (v. 3.7.5).

49. Latorre SM, Reyes-Avila CS, Malmgren A, Win J, Kamoun S, Burbano HA. Differential loss of effector

genes in three recently expanded pandemic clonal lineages of the rice blast fungus. BMC Biol. 2020;18:88.

50. O'Hanlon SJ, Rieux A, Farrer RA, Rosa GM, Waldman B, Bataille A, et al. Recent Asian origin of chytrid fungi causing global amphibian declines. Science. 2018;360:621–7.

51. Dale AL, Feau N, Everhart SE, Dhillon B, Wong B, Sheppard J, et al. Mitotic Recombination and Rapid Genome Evolution in the Invasive Forest Pathogen. mBio. 2019;10.

52. Pierce-Ward NT, Botvinnik OB, Reiter TE, Irber L, and Brown CT. Amino Acid k-mers enable assembly- and alignment-free sequence analysis. https://bluegenes.github.io. 2022.

53. Genome notes. https://www.ncbi.nlm.nih.gov/datasets/docs/v2/data-processing/policies-annotation/genome-processing/genome_notes/. Accessed 23 Jul 2026.
